# Sequence-dependent shape and stiffness of DNA and RNA double helices: hexanucleotide scale and beyond

**DOI:** 10.1101/2025.02.27.640607

**Authors:** Pavlína Slavníková, Marek Cuker, Eva Matoušková, Ivan Čmelo, Marie Zgarbová, Petr Jurečka, Filip Lankaš

## Abstract

The structure and deformability of double-stranded DNA and RNA depend on the sequence of bases, affecting biological processes and nanostructure design. Despite intense research, the dependence is incompletely understood. Here we present mechanical properties of DNA and RNA duplexes inferred from atomic-resolution, explicit-solvent molecular dynamics (MD) simulations of 107 DNA and 107 RNA oligomers containing all hexanucleotide sequences. The sequence-specific parameters include structure and stiffness at the rigid base level, the width and stiffness of major and minor grooves, and global material constants such as stretch modulus, twist rigidity, or bending and twisting persistence lengths. We propose a simple model to predict sequence-dependent shape and harmonic stiffness for arbitrary sequence, validate it on an independent set of MD simulations for DNA and RNA duplexes containing all pentamers, and demonstrate its utility in various applications. The large amount of simulated data enabled us to study rare events, such as base-pair opening lifetimes, or flips of the RNA sugar pucker into the B domain and the related dynamics of the 2’-OH group. Together, this work provides a comprehensive sequence-specific description of DNA and RNA duplex mechanics, forming a baseline for further research and allowing for a broad range of applications.

## Introduction

Double-stranded DNA (dsDNA) carries genetic information in all cellular life. RNA in biological contexts can take up a variety of structures, but its double helical form (dsRNA) is a prominent motif (1). Since the early days of DNA crystallography (2), a variety of experimental results have indicated that the DNA three-dimensional shape and structural flexibility depend on the sequence of bases (3–7), and analogous experimental data have been accumulating also for the RNA double helix (8–12). Sequence-dependent shape and stiffness of the NA double helices affect their interaction with proteins (13,14) and small ligands, some of them potential therapeutic agents (15–18). The knowledge of sequence-specific structural information greatly improves the performance of algorithms aimed at predicting transcription factor affinities (19–24) or identifying promoter regions (25,26). DNA sequence-dependent mechanical properties play an important role in supercoiling (27), nucleosome positioning (28–30), and chromatin structure (31), while the exact shape and stiffness of RNA helices influence the structural dynamics and tertiary contacts in large complexes such as the ribosome (32,33) and other assemblies (11). Both dsDNA and dsRNA are also key elements to build artificial NA nanostructures (34–36). It is therefore important to understand how the three-dimensional structure and deformability of DNA and RNA double helices depend on their sequence of bases.

The sequence-dependent shape and stiffness of NA duplexes can be described at various levels of detail, as reviewed (32,37–43). A widely used approach is the one where bases are modelled as rigid objects. Their relative displacement and rotation are defined by a set of internal coordinates, namely the intra-base pair coordinates shear, stretch, stagger, buckle, propeller and opening, as well as inter-base pair (or step) coordinates shift, slide, rise, tilt, roll and twist (44–47). Initial descriptions of DNA in terms of independent base-pair steps (3,48) and independent base pairs (49) were soon challenged by the discovery of structural correlations between different pairs and steps along the helix (45,48). The nature and properties of these couplings have been investigated in numerous studies (50–56). Various models, with different structural detail and interaction range, both harmonic and multistate, have been proposed (57–61). Parameters of the models are typically deduced from unrestrained molecular dynamics (MD) simulations of DNA oligomers involving all tetrameric sequences. The tetramer-oriented MD simulations, initiated by the ABC consortium (62,63), have resulted in a set of rules relating DNA base-pair and step structure to the base sequence (64) as well as to the backbone conformation (65). As for the RNA double helices, MD-derived information on sequence-dependent mechanical properties at the global scale (12,66), and at the tetrameric level (67), has been emerging. However, there is no guarantee that the tetrameric level will capture all the important sequence context effects in DNA and RNA helices. Indeed, both experimental evidence (68) and MD simulations (69) suggest that, at least for some DNA sequence motifs, a broader context may play a role.

The next sequence scale to consider, therefore, are pentamers. Simulations of pentameric sequences, each in many different contexts, were reported in (19). The extensive sequence coverage was made possible by the simplified underlying physical model, describing the DNA duplex in the space of selected collective and internal variables and using an implicit solvent model (70,71). Based on the simulated data, tools have been designed to analyze structural profiles of transcription binding sites (19,21). The approach found a variety of applications (e.g. (72,73)) and prompted further research aimed at identifying shape motifs in genomes (23,74). Importantly, the pentameric data include not only the shape (in terms of rigid base coordinates and minor groove widths) but also the flexibility (standard deviations of the above). Flexibility of the flanking sequences may affect transcription factor affinity for the core sequence motif (75,76). In a major development, a deep learning approach has been proposed to estimate sequence context effects on DNA shape and flexibility beyond the simulated pentameric data (22).

Building on extensive atomic-resolution, explicit-solvent MD simulations of DNA duplexes at the tetrameric level (63,64,69) and on earlier multivariate DNA models (59,77), atomistic MD simulations of DNA oligomers covering all pentameric sequences were used to parameterize a non-local, multistate mechanical model (60). The model involves two consecutive base-pair steps (i.e. steps within a trimer) embedded in their pentameric context. To our knowledge, sequence-dependent shape and stiffness of RNA duplexes have not been studied at the pentameric level of description.

Useful as it may be, the pentameric scale is awkward. Firstly, pentamers do not contain a central step and this asymmetry has to be dealt with in some way (22). Secondly, the pentameric scale does allow one to study sequence-dependent effects on the minor groove width (which spans a minimum of five nucleotides), but not on the major groove width, where at least the hexanucleotide level is needed (78). Finally, DNA A-tracts, a key sequence motif, exhibit a cooperative structure which is only fully developed at the length of six base pairs (79,80), and the same may be true for other motifs such as GGGCCC (81).

Thus, the hexanucleotide (or hexameric) scale seems a natural choice. Data at the hexanucleotide scale would cover the major groove, the cooperative A-tract and possibly other peculiar motifs, and provide a much richer information source on sequence-dependent structure and dynamics, allowing to test models, pass to longer scales, and answer a broad range of questions of biological importance. On the other hand, hexamers are a challenge. There are 2080 unique hexameric sequences, compared to 512 unique pentamers and 136 tetramers. Thus, the amount of data to be simulated is extensive. For instance, the pentameric simulations reported in (22) cover some pentamers as many as 250 times, but they still do not cover all hexamers.

At a more global level beyond rigid bases, one may characterize NA oligomers or their parts (fragments) just by a handful of material constants such as stretching, bending and twisting rigidity, and the elastic couplings between them. The values can be deduced from thermal fluctuations of suitably chosen global coordinates in unrestrained MD simulations. Considering fragments of different length, one may study, among other things, also the length dependence of these material constants. This approach was introduced in (82) and was largely expanded in later studies (42,52,83–87). An alternative method consists in pulling the oligomer by external force (66,88,89). The analysis of structural fluctuations at different pulling forces then allows one to probe the force dependence of the elastic constants (90,91).

Sequence-averaged material constants for DNA and RNA duplexes are well established (see e.g. (92) for an overview), but the knowledge about their sequence dependence is limited. Sequence-dependent DNA bending rigidity and the closely related bending persistence length have been studied experimentally (5) and computationally (93), the exceptionally high stretch modulus of DNA containing A-tracts has been experimentally established (6), and an early study reported sequence-dependent DNA twist-stretch (TS) coupling (94). Computational studies often focus on a limited range of sequences, such as those containing polypurines and dinucleotide repeats (66,80,82,89). A major experimental study established the cyclizability (i.e., the probability to circularize) for a large pool of DNA sequences (7), and subsequent work aimed at deducing sequence rules to predict this quantity (95,96). However, the cyclization probability depends in an intricate manner on both the shape and the stiffness of the duplex, a relation that is still a subject of intense research (e.g. (97–99) and references therein). Thus, a comprehensive understanding of sequence-dependent material properties for DNA and RNA duplexes is lacking.

One way to proceed is to directly probe these constants for a large pool of DNA and RNA duplex sequences. An attractive possibility, however, is also to establish sequence-dependent properties at the rigid base level and then consistently pass to the global level, as proposed by various authors (53,55,100–102). Here again the sequence-specific shape and stiffness at the hexameric scale may be of tremendous help.

In this work, we present atomic-resolution, explicit-solvent MD simulations and analysis of DNA and RNA duplexes involving all hexanucleotide sequences, compacted into a minimal set of 107 DNA and 107 RNA oligomers, each 33 base pairs (bp) long. Based on the MD data, we identify two length-scales of effective base-base interactions in DNA and RNA double helices. We then describe hexameric context effects on base-pair step and major groove structure and flexibility, and heptameric context effects on base-pair and minor groove structure and flexibility, where the heptameric properties are estimated from overlapping hexamers. Furthermore, we propose a simple non-local model of sequence-specific shape and harmonic deformability and parameterize it from the hexameric data. To validate the model, we produced an independent MD simulation set of 52 DNA and 52 RNA oligomers involving all pentameric sequences. The validation indicates excellent predictive power of the model, further corroborated by the specific model applications presented.

We then proceed to a more global level where the DNA or RNA oligomer as a whole, or its parts (fragments), are characterized by a small number of material constants. We estimate their values from thermal fluctuations of suitable global coordinates observed in our unrestrained MD simulations. The hexameric sequence variability enabled us to study not only the sequence-averaged values, but also their sequence-specific distribution and length dependence, greatly extending the limited information available so far. Furthermore, we examine the static disorder in DNA and RNA duplexes in terms of bending and twisting. Finally, the sheer amount of the simulated data allowed us to detect rare events, such as transient flips of the A-RNA sugar pucker into the B domain and the related motion of the 2’-OH group.

## Methods

### Sequence design

We designed a 516 base-pair (bp) minimal sequence containing each unique pentamer (512 in total) exactly once. To obtain the minimal sequence, a basic depth-first search approach was developed and implemented in-house, in the form of a Python script. While there are already several available tools for *k*-mer sequence generation (103), adding a simple, easily modifiable function was a matter of flexibility and convenience. The script, after being given a seed sequence (e.g. AAAAA), attempts to extend the sequence by another base. If the appended base forms a new *k*-mer, the addition is accepted and the script advances one base further. If the appended base does not form a new *k*-mer, the script attempts to change it for another base, and repeats the test. If the script exhausted all bases to no avail, the script backs up one base, changes it and repeats the test, thus in effect backing out of the unproductive branch. The resulting minimal sequence for pentamers was obtained within seconds on a desktop PC. The script is available at https://zenodo.org/records/14936465.

The script was also tested for other *k*-mer sizes, from dimers to nonamers. For odd-numbered *k*-mers, the developed depth-first search method seems to perform well, quickly producing optimal sequences.

However, this approach performed significantly worse for even-numbered *k*-mers, likely due to their possible self-complementarity, resulting in a less favourable search tree topology. Thus, to obtain the hexameric sequence, the ShortCAKE tool by Orenstein and Shamir (103) was used, as it provides optimal-length sequences for even-numbered *k*-mers. The optimal sequence containing all hexamers is 2145 bp long.

The optimal hexameric sequence was partitioned into 107 oligomers (set107), each 25 bp long (the ends were extended so as not to lose any hexamer upon the cutting), the minimal pentameric sequence was split into 52 oligomers, each 14 bp long (set52). Each sequence was capped by a GCGC tetramer at the ends to isolate possible end fraying (104). The same sequence set (with thymines mutated to uracils) was used for the RNA duplexes. In set107, the overwhelming majority of hexamers (3912 out of 4096) are present exactly once, and 184 exactly twice.

Similarly, the minimal pentameric sequence was partitioned into 52 DNA and 52 RNA oligomers (set52), each 14 bp long and capped by GCGC at each end. This set was used to validate the sequence-dependent shape and harmonic stiffness model we propose. The list of set52 sequences is in Tables S4 and S5.

To optimize the (single) hyperparameter of the model, namely the eigenvalue cutoff, a set of 14 oligomers (set14) covering all tetrameric sequences, each 22 bp long (14 bp plus the GCGC caps), was used. The list of s14 sequences is in Table S6. The MD data for the DNA version was taken from an earlier publication (65), the RNA version was simulated for this study. To examine the effective range of base-base interactions in DNA and RNA double helices, we also employed the MD data for a 33-bp oligomer in its DNA and RNA version from our previous work (87), here denoted by s0. Additionally, the DNA and RNA forms of s0 with neutralized phosphates and no ions were simulated.

### MD simulations and analysis

Atomic-resolution, explicit-solvent, unrestrained MD simulations were performed using the Amber 22 suite of programs. Each oligomer was built in its B-DNA or A-RNA form by the *nab* module of Amber and solvated with SPC/E waters in a truncated octahedron periodic box. K^+^ counterions were added to compensate for the NA charge, additional K^+^ and Cl^-^ ions were included to mimic the physiological concentration of 150 mM KCl. The OL15 force field (105) was used for the DNA oligomers. This force field, together with parmbsc1 (106) and more recent parameterizations tumuc1 (107), OL21 (108) and OL24 (109), belongs to the family of recent Amber force fields, and is considered a reliable representation of B-DNA oligomers (110–112). The well-established χOL3 force field (113–115) was employed for RNA, the ions were parameterized according to Dang (116). To produce the DNA and RNA neutral oligomers of s0, the +1 charge was split equally over the O1P and O2P oxygens and no ions were added. Each system was equilibrated by the standard ABC protocol (63), followed by a production run of 2 µs for each DNA sequence from the hexameric set107, and 1 µs for all the other systems, at 300 K and 1 atm. Hydrogen mass repartitioning (117) and a 4 fs time step were used for the production, snapshots were recorded every 10 ps. The *cpptraj* module of Amber (118) was employed to extract intra-base pair coordinates (shear, stretch, stagger, buckle propeller and opening), as well as base-pair step (shift, slide, rise, tilt, roll, twist) and helical coordinates (X-disp, Y-disp, helical rise, tip, inclination and helical twist) as defined by the 3DNA algorithm (46). Watson-Crick hydrogen bond lengths, and distances between phosphorus atoms used to define the groove widths, were also extracted with *cpptraj*. Altogether, this study covers all-atom, explicit-solvent, unrestrained MD simulations of 348 NA oligomers, the total simulation time is 0.45 ms.

Snapshots with at least one hydrogen bond broken within the central part (25 or 14 bp plus the flanking inner GC, cutoff 4 Å) were excluded from the analysis. We also excluded isolated snapshots with either all numbers in a *cpptraj* output line equal to zero, or with a negative twist, likely produced due to a numerical instability. Finally, we manually checked the plots of all the coordinates and excluded the (very few) parts with obvious structural anomalies, corresponding to non-canonical structures and discussed below. In this way, at least 89 % of snapshots in a DNA trajectory and 67 % in an RNA trajectory were left for further processing. Of note, three of the RNA sequences exhibited a H-bond break within the first 100 ns which was not repaired until the end of the simulation. These sequences were re-simulated using a different initial velocity seed, and the long-living break never appeared again.

We define the minor groove width as the distance between the phosphorus (P) atoms at the 3’ ends of a pentamer and assign it to the pentamer’s central pair (Fig. 1C). This definition is based on (78) where the authors proposed to take the mean of two neighbouring P-P distances, to be able to assign a minor groove width to a base-pair step, a definition implemented in 3DNA (46). However, taking just one distance suits better our purposes, since both of the P atoms are now inside a pentamer. Moreover, the definition employed here avoids artificial correlations between neighbouring minor groove widths, which occurs in the mean-distance definition just because two neighbouring widths share a common P-P distance. The major groove width is defined by the P-P distance within a hexamer and assigned to its central step (Fig. 1D), as in (46,78).

**Figure 1.**
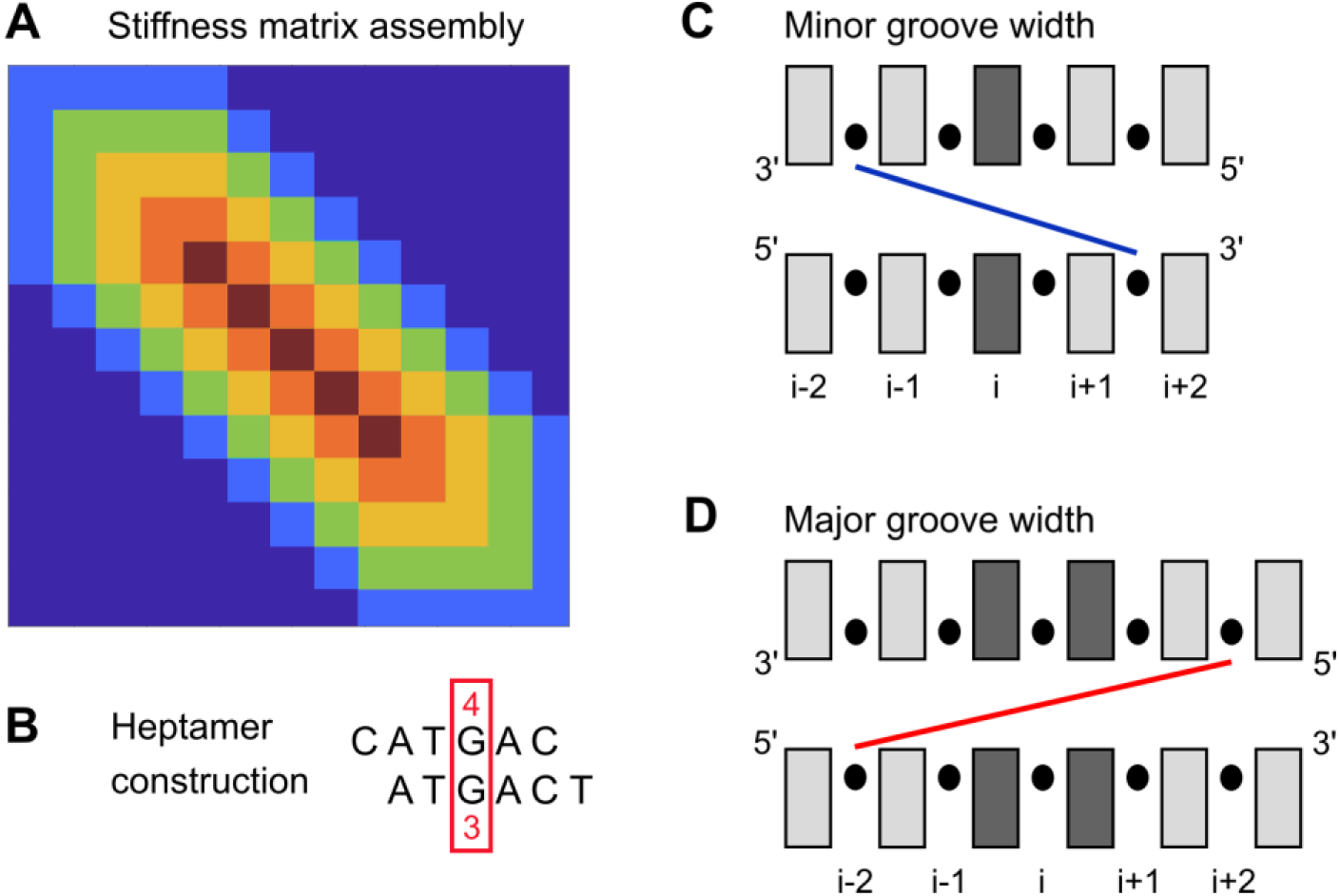
Constructions used in this work. (A) An optimal banded estimate of the stiffness matrix is first computed for each of the simulated 107 dsDNA and 107 dsRNA oligomers containing all unique hexamers. The hexameric blocks are then cut out of these matrices. Blocks for complementary hexamers are obtained by a simple transformation (see text). To create a stiffness matrix for an arbitrary sequence, the corresponding hexameric blocks are assembled as shown, with arithmetically averaged entries in the overlapping parts. The blocks exhibit multiple overlaps (zero to five marked by cyan, green, orange, red and dark red, respectively). The assembled matrix is zero outside the hexameric band (dark blue). (B) A heptamer constructed from overlapping hexamers. In this example, the heptamer CATGACT consists of two hexamers, CATGAC and ATGACT. The parameters for the heptamer’s central pair are computed as arithmetic means of the values in the two hexamers (4^th^ and 3^rd^ pair, respectively). (C) The minor groove width is defined as the distance between phosphorus atom centres at the 3’ sides of a pentamer in its heptameric context, and is assigned to the central pair. (D) The major groove width is the distance across a hexamer, and is assigned to its central step. To obtain the “empty” distances, 2 × 2.9 Å, or twice the phosphorus diameter, are to be subtracted.

### Deformation energy and stiffness

The free energy cost of distorting the oligomer, or deformation energy *E*, at the rigid base level is assumed in this work to be a quadratic function of the coordinate vector *w*,

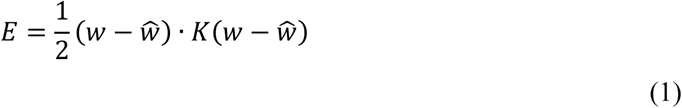

The equilibrium, minimum-energy, or ground-state coordinate vector *Ŵ* was estimated as the mean of *w* over the (filtered) MD trajectory, the stiffness matrix *K* was computed from the MD-derived coordinate covariance matrix Σ as *K* = *k*_*B*_*T*Σ^−1^.

The entries of the stiffness matrix are of three different physical dimensions depending on the combination of coordinates pertinent to the given entry: kcal/mol/Å^2^, kcal/mol/deg^2^ or kcal/mol/deg.Å (Eq. 1). If we want to study properties of the stiffness matrix as a whole, such as its eigenvalues and eigenvectors (as we do later in this work), the matrix entries have to be made dimensionally uniform, or dimensionless. We choose the length scale of 1 Å. As for the angle scale, the value numerically equal to the twist density of the B-DNA or A-RNA helix is a standard choice in extensible elastic rod descriptions (e.g. (82,119,120)). Taking the DNA value of 10.5 bp/turn (5,121,122) and its 3.23 Å inter-bp distance (123), we obtain the helical pitch of 10.5 × 3.23 = 34 Å and the twist density 360/34 = 10.6 deg/Å, so that the angle scale for DNA would be 10.6°. For the RNA helix we have 11.3 bp/turn (124) and 2.8 Å inter-bp distance (125), yielding the pitch of 31.6 ≈ 32 Å and the RNA angle scale 360/32 = 11.3°. Based on another way of reasoning, ref. (126) proposed 1/5 rad, or 11.5°. Here, to make things simple, we choose 11° as the angle scale both for DNA and RNA.

We also examine one-dimensional stiffness (or flexibility), i.e. the stiffness constant *k*_*x*_ for deforming the coordinate *x* (intra-base pair or step coordinate, major or minor groove width) while all the other degrees of freedom in the duplex are unconstrained. It is computed as *k*_*x*_ = *k*_*B*_*T*⁄var (*x*), where the variance is estimated from the time series of *x* extracted from the MD trajectory.

### Estimating sequence-dependent shape

To define the equilibrium conformation of a base-pair step, we take the average coordinates over the MD trajectory for the step in its hexameric context. The major groove width is inferred for the given hexamer in a similar fashion and assigned to the central step. From the unique hexamers, values for their complements are constructed, transforming the coordinates of the central step according to their parity: odd coordinates (shift, tilt, Y-disp, tip) change sign, the other, even ones (including major groove width) remain unchanged. Since the hexameric sequence dependence is assumed, the odd coordinates of the central step of self-complementary hexamers (e.g. ATATAT) are set to zero.

To define the equilibrium structure of a given pair or minor groove width, we construct a heptamer by considering two overlapping hexamers, and compute the arithmetic mean of the values for the pair in the centre (Fig. 1B). Coordinate values for complementary heptamers are then constructed. The odd coordinates (shear, buckle) change sign, the even ones (including minor groove width) remain unchanged. The shape of an arbitrary sequence is then estimated by applying a hexameric/heptameric sliding window (thus, three pairs at each end are not covered).

### Estimating sequence-dependent stiffness

To extract hexameric stiffness blocks from our MD data, we first compute, for the inner 25-bp part of each oligomer in set107, a stiffness matrix that is exactly zero outside a band defined by the overlapping hexameric blocks (Fig. 1A). To this end, we use the maximum absolute entropy approach introduced in (127). By construction, the covariance related to this banded stiffness is equal to the original covariance matrix within the band (but may be different outside the band). The case considered in (127) only involved blocks with a single overlap, whereas our blocks exhibit up to five overlaps (Fig. 1A). Nevertheless, the method applies to multiple overlaps as well (128) and is as follows. Each of the diagonal hexameric blocks of the original covariance matrix is inverted and written to the corresponding block in the stiffness matrix, with all entries in the overlapping regions summed up. Then the covariance blocks corresponding to overlaps between adjacent hexamers are inverted and subtracted from the corresponding blocks in the stiffness matrix. Finally, the stiffness matrix is multiplied by *k*_*B*_*T*.

We apply this procedure to the simulated 107 dsDNA and 107 dsRNA oligomers to obtain the banded estimates of their stiffness matrices, then we cut out the hexameric blocks from them. To obtain stiffness blocks for complementary pentamers, the entries have to be rearranged (the last pair is now the first one etc.) Moreover, the entries corresponding to odd-even combinations (e.g. tilt-twist) change sign (57). To estimate the stiffness matrix of an arbitrary sequence, we assemble it from the hexameric blocks (Fig. 1A). The overlapping parts are arithmetic means of the individual blocks. Notice that, by construction, the banded stiffness matrices of the sequences from which the blocks were cut out (the majority of the 107 DNA and 107 RNA sequences, save possibly for the very few duplicate hexamers), are reproduced exactly. The arithmetic averaging is certainly not the only possible: the averaged blocks are positive definite and various methods for averaging positive definite matrices have been proposed (129). We also tested the harmonic average (i.e. arithmetic average of the inverses), with nearly identical results.

## Results and discussion

### Two length scales of effective base-base interactions

In many earlier studies, the range of interactions between rigid bases and base pairs to be included in the proposed model was chosen *a priori*. Here we start by asking a simple question: what is the effective interaction range between bases in DNA and RNA double helices? To this end, we first analysed stiffness matrices for the 33-bp oligomer in its DNA and RNA versions from earlier work (87) (here denoted sequence s0). Our approach is illustrated in Fig 2.

**Figure 2.**
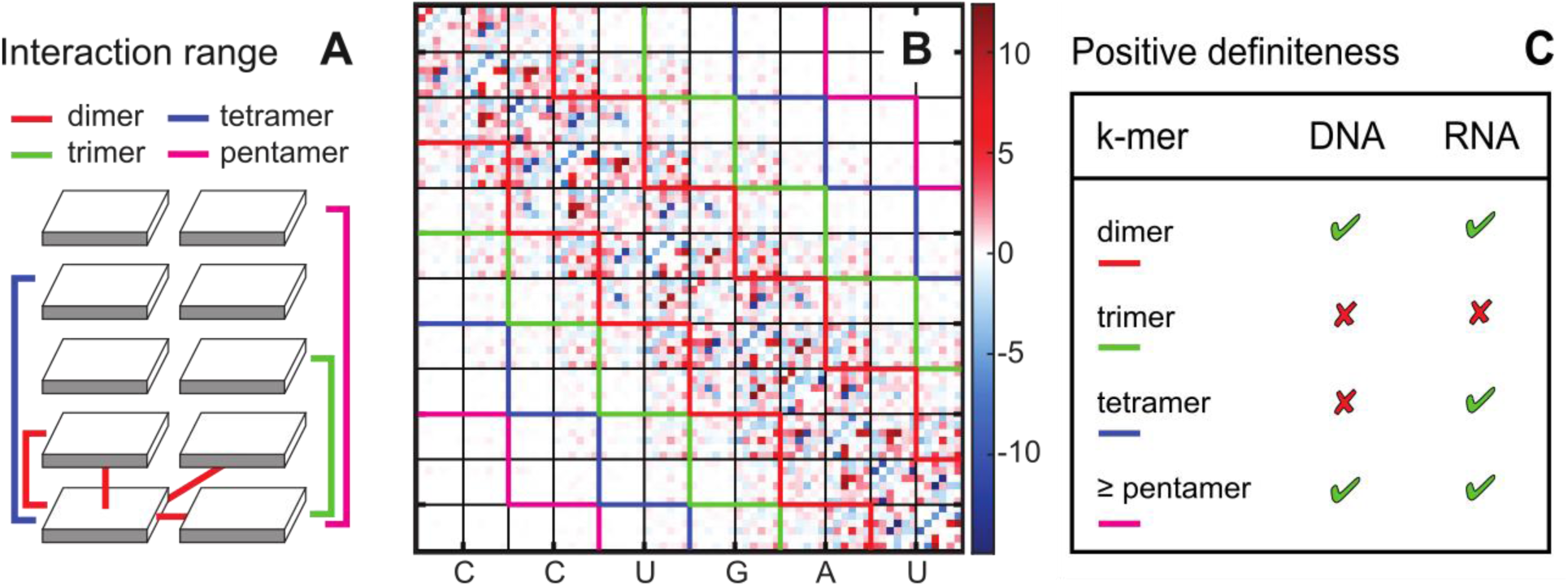
Effective base-base interactions in a double helix. (A) The nearest-neighbour, or dimeric interactions (red) comprise hydrogen bonding and intra-strand as well as inter-strand stacking. If these were the only interactions in the helix, the stiffness matrix (B) would be exactly zero outside the red-delineated band (shown here is the central part for the s0 sequence in its RNA version). Similarly, if interactions were limited to trimers (green) or tetramers (blue), then again the stiffness matrix would be zero outside the corresponding band, which is not the case. (C) If the trimeric range is imposed (i.e. the entries outside the band are set to zero), the resulting matrix is not positive definite. Thus, important long-range interactions are missed in this way. It is only the pentameric range (magenta) or longer that produce a positive definite matrix both for DNA and RNA helices. This result holds for the s0 sequence as well as for the simulated 107 DNA and 107 RNA oligomers containing all hexanucleotides. Moreover, the result also holds for the s0 sequence in which phosphate charges were neutralized and no ions present in the simulation. This indicates that the long-range base-base interactions are not dominated by electrostatic effects.

First of all, bases in DNA or RNA double helices are involved in nearest-neighbour, or dimeric, interactions of Watson-Crick hydrogen bonding and intra-strand as well as inter-strand stacking (red connecting lines in Fig. 2A), which have been studied in great detail (130–133). If these were the only interactions present, the entries of the stiffness matrix outside the dimeric range (delineated by red contours in Fig. 2B) would be exactly zero, as discussed (50). This is clearly not the case, as there are visibly non-zero entries outside this range. If the dimeric range is imposed by setting all entries outside the red band to zero, the resulting matrix remains positive definite, albeit with eigenvalues far away from those of the full matrix (Figs S1, S2). One might think that adding the trimeric interaction range (green lines in Fig. 2A, B) could only improve matters. However, the opposite is true – the resulting matrix is not positive definite and therefore cannot represent a valid stiffness matrix. Thus, important longer-range interactions are missed. Adding the tetrameric interaction range (blue in Fig. 2A, B) produces a positive definite matrix for RNA, but not for DNA, and the eigenvalues are still quite off (Figs S1, S2). It is only the pentameric interaction range (magenta in Fig. 2A, B) or longer that again produce a safely positive definite matrix both for dsDNA and dsRNA. Motivated by these observations, we tested all the simulated DNA and RNA oligomers (set107, set52 and set14), with entirely analogous results.

The most likely contributors to DNA stiffness are base stacking interactions and electrostatic repulsions between charged phosphates, although their exact roles are still debated (134,135). Stacking interactions are short-range, but charge repulsion could in principle contribute to long-range couplings. Bending persistence length decreases with increasing salt concentration, while the stretch modulus increases, both for dsDNA and dsRNA (136,137), and DNA twist rigidity appears to be independent of salt (138). A useful method to probe the electrostatic effects is a partial or complete neutralization of the phosphate charges (134,139,140). The bending persistence length of dsRNA is higher than that of dsDNA, but a recent study reported that this difference decreases with increasing monovalent salt, and the two become identical in simulated oligomers with neutral phosphates (141) (the authors dispersed the +1 charge over the whole phosphate group rather than just over the O1P and O2P oxygens).

To test the role of electrostatic interactions in long-range base-base elastic couplings, we simulated the s0 sequence in its DNA and RNA forms with the phosphate charges neutralized (Methods) and no ions present. The results were very similar to the charged version (Figs S1, S2). Together, these observations indicate that at least the pentameric scale, i.e. roughly half the helical turn, is necessary to capture the effective base-base interactions in DNA and RNA double helices, and that these couplings are not dominated by electrostatic interactions. It is possible that the couplings stem from the mechanical constraints imposed by the backbone, a phenomenon known from polymer physics (142) and considered also in DNA modelling (54).

### Sequence-averaged DNA and RNA double helices

We performed atomic-resolution, explicit-solvent MD simulations of 107 dsDNA and 107 dsRNA oligomers containing all hexanucleotide (or hexameric) sequences, the vast majority of them exactly once. The oligomers are 33 bp long and involve the hexamer-containing central 25 bp part capped by the GCGC sequence at each end. Prior to any further processing, the data were filtered for broken pairs, non-canonical structures, and numerical anomalies (Methods).

From the filtered MD data, we extracted the trajectory means of the step coordinates for the central steps of individual hexamers, the hexamer-specific major groove widths, as well as intra-base pair coordinates for the central pairs of individual heptamers, and the minor groove widths for the central pentamer of individual heptamers. The heptameric values were estimated from overlapping hexamers (Fig. 1B). Besides the means (first moment), we also obtained the variances (second moment) and used them to compute flexibilities in terms of the effective stiffness constants *k*_*x*_ (Methods).

Since the sequences include all unique hexamers, nearly all of them exactly once, the simulated data represent a comprehensive, well-balanced sequence set. To obtain sequence-averaged values, we average the step coordinates and major groove widths over all hexamers, and intra-base pair coordinates and minor groove widths over all heptamers. One strand has to be chosen as a reference. Upon changing the reference strand, the odd coordinates (shear, buckle, tilt, shift, tip and Y-disp) change sign, while all the other, even ones remain unchanged. Since there is no physical reason to choose one strand or the other, we can imagine doing the averaging such that one strand and subsequently the other one is chosen as a reference. Consequently, the means of the odd coordinates are exactly zero.

Table 1 shows the sequence-averaged values, each with the standard deviation indicating the spread over individual hexamers or heptamers. The values conform to the B-DNA or A-RNA structures. The sequence-dependent variability may be quite substantial, e.g. for roll, twist or inclination. The variability is in general smaller for RNA than for DNA, indicating that the shape of the A-RNA helix is on average less modulated by the sequence than in the B-DNA case. A notable exception is inclination and especially roll, which are more variable in RNA helices.

**Table 1.** Sequence-averaged structural parameters of the simulated DNA and RNA helices. The standard deviations represent sequence-specific variability over all hexameric or heptameric sequences.

|  | DNA | RNA |
| --- | --- | --- |
| Shear [Å] | $0 \pm 0.12$ | $0 \pm 0.12$ |
| Stretch [Å] | $-0.02 \pm 0.04$ | $-0.02 \pm 0.05$ |
| Stagger [Å] | $0.03 \pm 0.10$ | $-0.07 \pm 0.10$ |
| Buckle [°] | $0 \pm 5.86$ | $0 \pm 4.79$ |
| Propeller [°] | $-11.03 \pm 3.70$ | $-12.61 \pm 1.75$ |
| Opening [°] | $0.24 \pm 0.84$ | $0.28 \pm 0.83$ |
| Shift [Å] | $0 \pm 0.33$ | $0 \pm 0.09$ |
| Slide [Å] | $-0.13 \pm 0.36$ | $-1.64 \pm 0.14$ |
| Rise [Å] | $3.33 \pm 0.10$ | $3.30 \pm 0.10$ |
| Tilt [°] | $0 \pm 1.61$ | $0 \pm 0.72$ |
| Roll [°] | $3.14 \pm 2.53$ | $8.83 \pm 2.95$ |
| Twist [°] | $34.49 \pm 2.84$ | $30.08 \pm 1.01$ |
| X-disp [Å] | $-0.84 \pm 0.53$ | $-4.42 \pm 0.31$ |
| Y-disp [Å] | $0 \pm 0.41$ | $0 \pm 0.17$ |
| h-Rise [Å] | $3.24 \pm 0.11$ | $2.67 \pm 0.14$ |
| Inclination [°] | $5.72 \pm 4.52$ | $15.93 \pm 4.63$ |
| Tip [°] | $0 \pm 2.62$ | $0 \pm 1.34$ |
| h-Twist [°] | $35.54 \pm 2.75$ | $32.24 \pm 1.64$ |
| Minor g. width [Å] | $12.04 \pm 0.89$ | $17.11 \pm 0.22$ |
| Major g. width [Å] | $18.36 \pm 0.61$ | $18.11 \pm 0.31$ |

A number of studies compared the structure of MD-simulated DNA and RNA oligomers to crystallographic or NMR data (e.g. (32,64,110,111,115,143). Although the comparison may not be straightforward, the general conclusion is that, among other parameterizations, the DNA and RNA force fields employed here (OL15 for DNA (105), χOL3 for RNA (113,114)), and the simulation setup used, give a reliable structural prediction. Here we only briefly discuss two coordinates, rise and twist.

The DNA helical rise (h-rise, Table 1), or distance between base-pair centres projected onto the helical axis, agrees quantitatively with recent measurements of base-pair distance using gold labels and anomalous small-angle x-ray scattering (3.23 ± 0.1 Å) (123), while the RNA h-rise is somewhat underestimated compared to the consensus experimental value of 2.8 ± 0.1 Å (125). The local rise, capturing the distance between successive base pairs along their mean normal, primarily reflects the local stacking interactions and is very similar in the DNA and RNA helices.

Twist is an interesting case. The MD-derived helical twist (h-twist, Table 1), or the twist angle about the helical axis, implies 11.2 bp/turn for RNA, in agreement with the experimental value of 11.3 ± 0.1 bp/turn (124), while the local RNA twist is far from that value. Indeed, local and helical twists are expected to differ due to the A-like structure of the RNA helix. In the case of DNA, however, it is the local twist that yields 10.5 bp/turn, a value predicted for a random sequence with 50 % GC content based on DNA cyclization (5) and in agreement with 10.4 ± 0.1 bp/turn from gel electrophoresis (121) and 10.6 ± 0.1 bp/turn from enzyme digestion (122). The DNA helical twist, in contrast, is ∼1° higher than the local one and gives just 10.2 bp/turn, although the two types of twist are generally assumed to be similar for B-DNA. This issue may deserve further investigation.

### Sequence-dependent structure and flexibility

The complete hexanucleotide coverage enables us to examine the sequence dependence at an unprecedented context range with minimal model assumptions. However, visualizing the data is not straightforward. The way we proceed is to present the values for central steps in individual tetramers, and central pairs within pentamers, each averaged over all its hexameric (heptameric) contexts. We then separately present the hexameric/heptameric variability in terms of the difference between the highest and the lowest value for the given central tetramer/pentamer. As for the flexibility, we quantify the context dependence in relative terms, i.e. maximal minus minimal value divided by the average. Fig. 3 illustrates the information we provide. We find this type of visualization (a “heatmap”) useful, since it enables the reader to discern sequence-dependent patterns at a glance.

**Figure 3.**
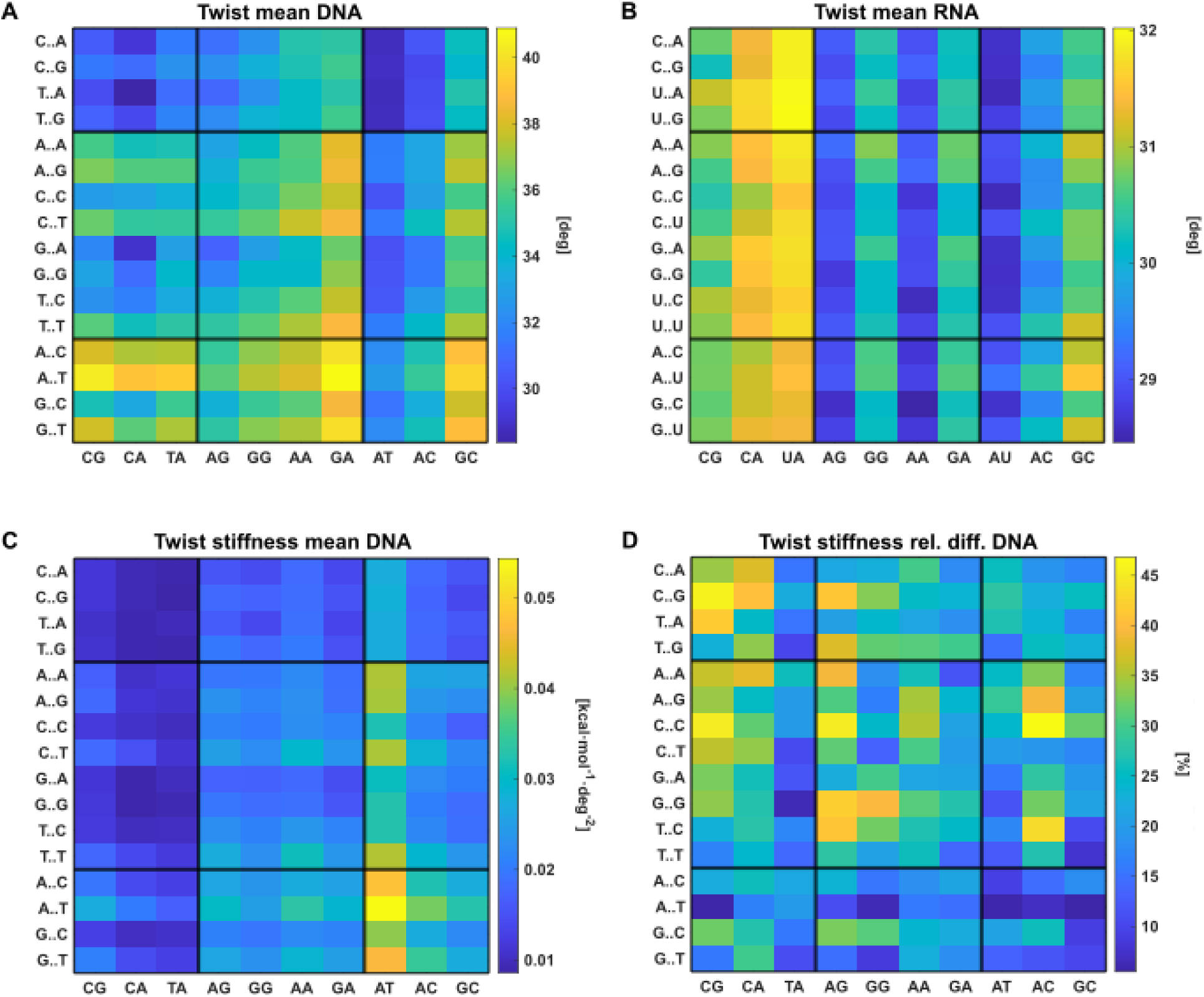
(A-B) DNA and RNA twist of the central step of the tetrameric sequence indicated, averaged over all the hexameric contexts. Sequence-specific rules, quite different for DNA and RNA duplexes, can be seen. (C) Tetramer-specific twist stiffness reveals the extreme rigidity of the AATT sequence in all hexanucleotide contexts. (D) The relative variability of DNA twist stiffness over the hexameric contexts of the given tetramer reaches up to 45 %. The black lines separate the YR, RR, and RY sequences (R – purine, Y – pyrimidine). This type of visualization (a “heatmap”) enables one to quickly discern sequence-dependent patterns in the data, such as domains of high and low values or the degree to which the values are already determined by the central step. Analogous plots for the remaining coordinates are at https://zenodo.org/records/14936465.

Taking twist as an example (Fig. 3), we see that the range of DNA values is much broader than for RNA. We can discern the domain of low (the YYRR tetramers, upper left corner of Fig. 3A) and high DNA twist (RYRY, lower left corner). In some cases, the values are largely dictated by the central step (DNA AT and most RNA steps). However, on can also discern hexanucleotide context effects undetectable in earlier tetrameric data (64,67). For instance, the hexamer-context dependent stiffness variability can reach up to 45 %, is on average highest for YYRR (Fig. 3D, upper left corner) and lowest for RRYY (lower right corner).

Analogous plots for the remaining coordinates are at https://zenodo.org/records/14936465. In particular, it is observed that, with the exception of tilt, the hexameric stiffness variability for RNA step coordinates is roughly half the DNA value. We also point out the extreme flexibility of DNA and RNA buckle (< 0.01 kcal/mol/deg^2^).

The sequence-specific minor groove widths are shown in Fig. 4. Here again the spread of DNA values is much larger than that of RNA. While the DNA minor groove width is mostly determined by the outer bases of the pentamer (Y…R > R…R > R…Y on average), the RNA values are largely dictated by the central trimer, or even by the central pair (A-U or G-C). Moreover, the RNA minor groove is much stiffer than the DNA one and its heptameric variability is smaller. As for the major groove, its width and stiffness depend on the whole hexameric sequence and we only present their histograms (Figs S3, S4). Tables in the csv format of means and stiffnesses for all the individual hexamers or heptamers can be found at https://zenodo.org/records/14936465.

**Figure 4.**
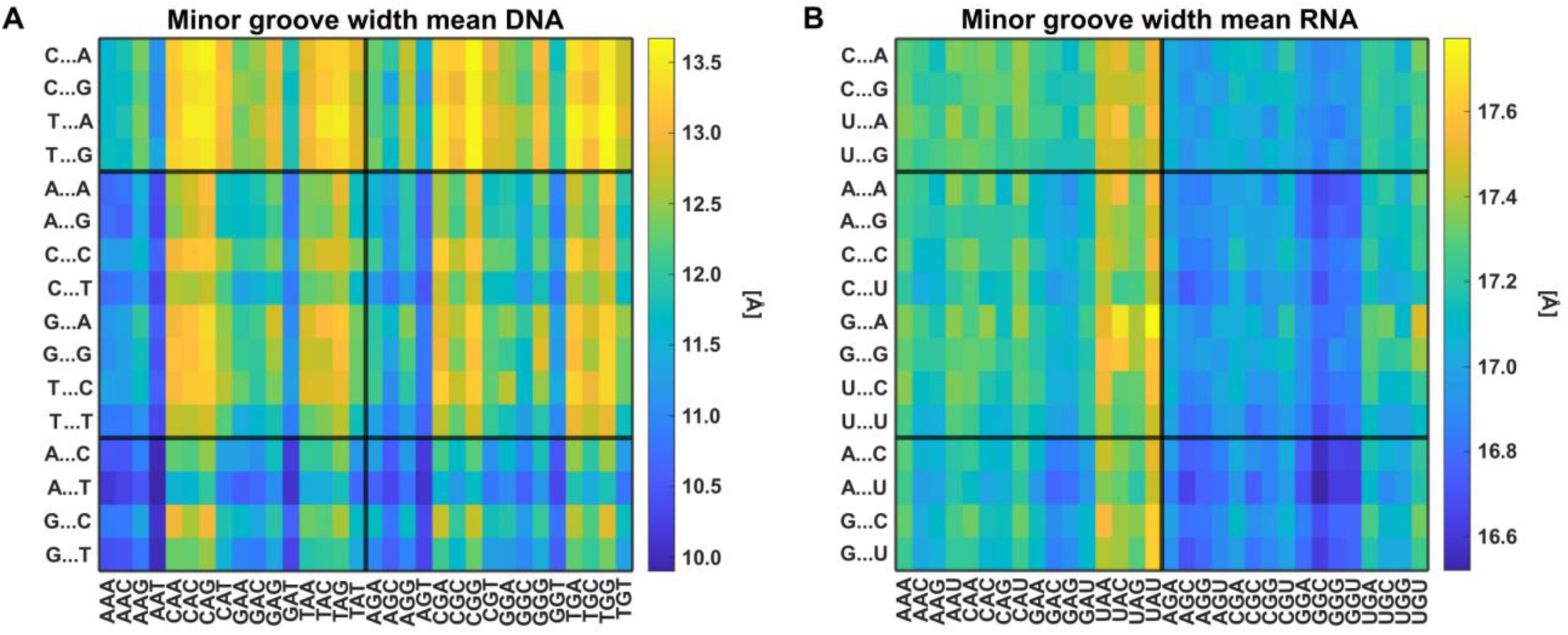
Sequence-dependent variability of the minor groove width. Values for pentameric sequences, each averaged over all its heptameric contexts, for dsDNA (A) and dsRNA (B). Notice the contrasting patterns of sequence dependence and the smaller spread of values for the RNA double helices.

### Global elasticity

At the global level, we characterize the simulated 107 DNA and 107 RNA duplexes in terms of their bending and twisting persistence lengths, stretch modulus, twist stiffness and the coupling between twisting and stretching (TS coupling). The large, balanced set of sequences containing all hexamers, most of them exactly once, enables us to examine sequence variability of these constants, greatly extending the available information.

To compute the bending persistence length (p.l.) of a given oligomer, we probe directional correlations between its base-pair normals. The dynamic p.l. *l*_*d*_ is obtained by factoring out the static structure as proposed in (93), with an additional prefactor *α*_*d*_. It is defined by the relation

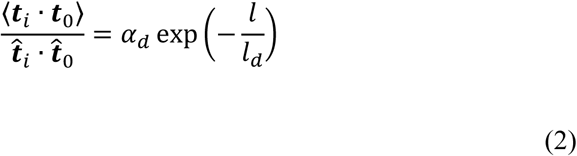

where ***t***_*i*_ and ***t***_0_ are instantaneous normals of base pairs 0 and *i* separated by the distance *l*, ***t̂***_*i*_ and ***t̂***_0_ are normals of the same pairs in the equilibrium structure (reconstructed from hexamer-specific values of base-pair step coordinates), the brackets denote mean over the (filtered) trajectory, and *l* is the trajectory mean of the sum of helical rises between the two pairs. The values of *l*_*d*_ and *α*_*d*_ are deduced from fitting a straight line to the logarithm of the left-hand side as a function of *l* (semilog plot). The prefactor *α*_*d*_ accounts for the possibility that the fitting line does not pass through the origin (42,144).

The twist p.l. *l*_*tw*_ probed here reflects the correlation length between helical twists, includes again a constant prefactor (42), and is defined by

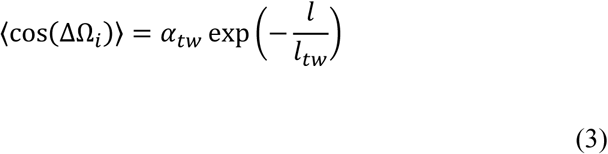

where Ω_*i*_ is the sum of helical twists between pairs 0 and *i*, and ΔΩ_*i*_ = Ω_*i*_ − 〈Ω_*i*_〉. Again, a linear fit of the semilog plot is used to deduce *l*_*tw*_ and *α*_*tw*_. Since the mean twist is subtracted, the twist p.l. is analogous to the dynamic bending p.l.

To infer the stretch and twist moduli and the TS coupling, we examine thermal fluctuations of the total length and twist of the oligomer, as introduced in (82). These global coordinates have to be carefully chosen. Here we define the oligomer length as the sum of helical rises, and its global twist as the sum of helical twists, using their 3DNA definitions (46). This choice, proposed in (83), gives the correct magnitude and sign of the TS couplings for DNA and RNA (83), realistic values of stretch and twist moduli for DNA (42) as well as RNA and hybrid duplexes (87), and has been used in many subsequent studies.

Table 2 summarizes the mean values and standard deviations of the material constants for the set of 107 DNA and 107 RNA sequences, their distributions are in Figs S5 and S6. For each oligomer, the central 25 bp part was examined, the GCGC caps were excluded. The stretch and twist moduli were deduced from the fluctuations of the corresponding global coordinate while the other one was neglected (integrated out), the TS coupling was inferred from the 2-by-2 stiffness matrix involving both coordinates, as detailed in (42).

**Table 2.** Global material constants for the set of DNA and RNA oligomers containing all hexanucleotide sequences.

| Parameter | Symbol | DNA<sup>a</sup> | RNA<sup>a</sup> |
| --- | --- | --- | --- |
| Stretch modulus (pN) | $Y$ | $1621 \pm 127$<br>*900 - 1500* | $694 \pm 37$<br>*350 - 500* |
| Twist stiffness (nm) | $C$ | $119 \pm 7$<br>*94 - 109* | $118 \pm 9$<br>*100 \pm 2* |
| Half twist persistence length (nm) | $l_{tw}/2$ | $136 \pm 8$ | $122 \pm 11$ |
| Twist-stretch coupling (nm/turn) | $k$ | $0.81 \pm 0.10$<br>*0.42 - 0.5* | $-1.11 \pm 0.20$<br>*-0.85 \pm 0.4* |
| Dynamic bending persistence length (nm) | $l_d$ | $68 \pm 4$<br>*46 - 54, 78 \pm 13* | $84 \pm 4$ |

<sup>a</sup> Experimental values (*in italics*) compiled from the literature, see text.

The sequence-averaged values of the stretch modulus, twist stiffness and TS coupling reproduce well-known experimental findings: RNA is much more flexible in stretching than DNA, has a similar twist modulus, and its TS coupling is of similar magnitude and opposite sign. The values are somewhat overestimated compared to the experimental range (5,6,92,145), as already reported for a limited set of DNA and RNA sequences (42,87,91). However, here we also provide the sequence-specific variation over a large, well-balanced set of oligomers containing all hexanucleotides. Taking three standard deviations as the variability measure, we obtain a spread of roughly 20 % for *Y* and *C*, while the TS couplings may vary by as much as 50 %.

The dynamic bending p.l. *l*_*d*_ captures the decay length of directional correlations due to thermal fluctuations. Together with the static disorder quantified by the static p.l. *l*_*s*_, they make up the total p.l. *l*_*lot*_ through the relation 1/*l*_*lot*_ = 1⁄*l*_*d*_ + 1⁄*l*_*s*_ (146). Taking the DNA value of *l*_*d*_ = 78 *nm* from cryo-EM experiments (145), and the consensus value *l*_*lot*_ = 50 *nm*, implies *l*_*s*_ = 139 *nm*. In contrast, cyclization experiments indicate a very large *l*_*s*_ (> 1000 nm), so that *l*_*lot*_ = *l*_*d*_ with high precision (147), a discrepancy probably due to different experimental conditions (148). The large *l*_*s*_ is also supported by an earlier simulation study (93). If so, then the 46 – 54 nm experimental sequence-dependent range (taken from (5)) corresponds to *l*_*d*_ and the MD value of *l*_*d*_ is again somewhat overestimated. Its sequence-dependent variability of 18 % found here is consistent with the earlier computations (93) and comparable to the experimental one.

The experimental RNA total persistence length is higher than the DNA one in physiological conditions (92,125,141). We are unaware of any experimental value of RNA dynamic p.l., but our computed value is again higher than the DNA one, as in earlier simulation studies (87,126). Furthermore, our sequence-dependent variation indicates a similar spread of values as for the DNA dynamic persistence length.

The twist stiffness *C* and twist persistence length *l*_*tw*_ are often used interchangeably but are two different quantities, the former capturing the mechanical rigidity and the latter the decorrelation length of twist fluctuations. They are closely related for an intrinsically straight, homogeneous, isotropic elastic rod, where *C* = *l*_*tw*_⁄2 (derived e.g. in (149,150)). Here we find that for RNA the two indeed coincide, but our deduced *l*_*tw*_⁄2 is distinctly higher than *C* for DNA.

Early MD simulations already indicated that DNA elastic properties are length-dependent: short fragments of a given oligomer are more flexible, the rigidity increases with the fragment length, and it only gets saturated at the scale of half the helical turn or more (82). The phenomenon was later re-discovered using much longer oligomers and simulated time scales (52), and was a subject of intense theoretical study (51,53–55). Still, many questions remain open. Most studies focus on bending and twisting rather than stretching, and if stretching is included, irrelevant length definitions are sometimes used. As for RNA, the authors often consider local descriptions (local twist, roll, etc.) pertinent for the stripe of base pairs surrounding the central hollow, rather than for the helix as a whole. The length-dependence of TS coupling, to our knowledge, has not been examined. Moreover, a limited set of sequences have so far been probed.

Here we study the length-dependent elasticity of DNA and RNA duplexes on our sequence set covering all hexanucleotides. First of all, we find that the bending and twisting persistence lengths are nearly length-scale independent. Indeed, we do not observe any obvious change of slope in our semilog plots used to infer the persistence lengths. In contrast, the global elastic constants depend significantly on the fragment length. Both the stretch modulus and the twist stiffness increase with length, and the increase ranges from some 30% for RNA stretch modulus up to a three-fold increase for DNA twist stiffness. TS couplings slowly decrease for longer fragments, but a non-monotonic behaviour is observed at shorter scales. Importantly, although the values are sequence-dependent, the shapes of all the curves appear to be universal, and the 25 bp scale examined here seems to be about the minimal length to obtain converged values.

### Bending and twisting static disorder

The static structural disorder in DNA and RNA double helices may be generated due to the sequence-dependent variability of the equilibrium (or static) structure. A single, short oligomer simply has a particular static structure and one cannot speak about a disorder. However, the structure of a very long stretch of the double helix with random sequence, or a large ensemble of shorter helices, may exhibit a static noise. It has long been known that the static disorder with respect to bending may be quantified by the static bending persistence length *l*_*s*_, defined by the relation 〈***t***_*i*_ · ***t***_0_〉 = exp(− *l*⁄*l*_*s*_), where the brackets now denote the average over the static structural ensemble (146). A disadvantage of this definition is that the sequence-averaged structure over which the static disorder is imposed may itself not be straight. This is the case especially for RNA helices where the A-form implies a significant inclination of the base pairs with respect to the helical axis, resulting in periodic changes of the angle between the base-pair normals with the helical repeat. To deal with this problem, we propose to compute the static bending p.l. from the definition entirely analogous to the dynamic one (Eq. 2), but now taking the static structural ensemble in place of the MD trajectory snapshots. In this way, the sequence-averaged static structure is factored out.

Just as the static disorder with respect to bending, one may study the disorder with respect to twisting which, we believe, has not been examined so far. We again use the definition entirely analogous to the twist (dynamic) persistence length (Eq. 3) to define the twist static persistence length, considering the static ensemble instead of the ensemble of MD snapshots.

We infer the static bending and twisting persistence lengths based on the hexanucleotide structural data. To do so, we generate 10^5^ random DNA sequences and the same number of random RNA sequences, each 500 bp long, and assign a static structure to each of them to obtain the structural ensemble, then use Eqs 2 and 3 to infer the persistence lengths. The periodic oscillations are nicely factored out, and the semilog plots are very close to linear (Fig. S7). We find that the static disorder, both with respect to bending and twisting and both for DNA and RNA, is very weak. The inferred DNA static bending p.l. is 963 nm and the RNA one equals to 890 nm, in line with the scenario proposed in (147). The twist disorder we found is even weaker, the DNA and RNA half static twist p.l. being 1647 and 3687 nm, respectively. Thus, the MD-derived dynamic persistence lengths in Table 2 should be also understood as the total ones (we are not aware of any relation for the twist p.l. analogous to 1⁄*l*_*lot*_ = 1⁄*l*_*d*_ + 1⁄*l*_*s*_, but it is reasonable to assume that the static and thermal noise superimpose in some way).

### A predictive model of sequence-dependent shape and stiffness

#### Shape

We estimate the static (equilibrium) structure, or shape, of an arbitrary sequence from the hexanucleotide data using a hexameric sliding window for base-pair step coordinates and major groove widths, and a heptameric sliding window for intra-base pair coordinates and minor groove widths. The heptameric data are computed as averages over the two overlapping hexamers forming the heptamer (Fig. 1B). The one-dimensional stiffness associated with the individual coordinates (Methods) is estimated analogously.

Previous studies used sliding windows of tetrameric (e.g. (59,64)) or pentameric length (22,60). The obvious limitation of a sliding window approach is that no context effects beyond the window length can be captured. At least two methods have been proposed to overcome this problem. A model has been introduced where the shape is related to the inverse of a narrowly banded, but not block-diagonal matrix (57,58,126). Since the inverse of a banded (but not block diagonal) matrix is in general dense, the (relatively few) parameters defining the banded matrix transform into a possibly long-range sequence dependence of the shape. Another method is based on a deep learning approach where the missing sequence contexts are predicted by the trained model (22), and indeed, the authors reported small but discernible effects beyond the pentameric scale (22).

Here, by contrast, we are limited by the window length. On the other hand, our sliding windows are longer than any used before, the approach is entirely straightforward (in contrast to the highly non-trivial model parameterizations in (57,58,126) or (22)), and involves no model assumptions other than the underlying all-atom MD data (and the heptamer approximation).

To test the predictive power of our model, we produced a validation set of independent all-atom MD simulations of 52 DNA and 52 RNA oligomers containing all pentameric sequences (set52, Methods). For each sequence, we predicted its shape from the hexameric/heptameric model and compared to the actual values from MD. The results are in Tables S1 and S2. The average error in DNA step coordinates is < 0.5° and < 0.05 Å, the RNA values are even smaller. The intra-base pair coordinates are also well predicted, with the exception of the very flexible DNA buckle where the error reaches nearly 1°. The sequence-dependent DNA minor groove width is captured within 0.16 Å on average, the major groove within 0.26 Å, the RNA values are again even better.

#### Stiffness

Just as for the shape parameters, we tested the prediction of the stiffness constants associated with individual coordinates (Methods). The relative errors *ε* = |*k*_*pred*_ − *k*_*set*52_|⁄*k*_*set*52_ are listed in Table S3. The DNA step stiffnesses are predicted within 3.5 % (twist 1.2 %), the intra-base pair ones within 5 %, the groove stiffnesses within 1.5 %. The RNA data are again considerably better.

Encouraged by these results, we set out to predict not only the shape and coordinate flexibility, but the whole stiffness matrix of an arbitrary sequence. The stiffness matrix is related to the full set of intra-base pair and step coordinates of the sequence and involves the diagonal terms as well as the elastic couplings between any two of them. This approach, introduced in (50), extends the traditional models of independent base-pair steps (3,48) and base pairs (49).

To construct the stiffness matrix for an arbitrary sequence, we took the hexameric stiffness blocks and assembled them together, arithmetically averaging the overlapping parts (Fig. 1A, Methods). We again used the set of 52 DNA and 52 RNA oligomers (set52) covering all pentamers to validate the model. To our disappointment, the smallest eigenvalues of some of the assembled matrices (non-dimensionalized as in Methods) for the set52 sequences were very close to zero, or even negative (Fig. S9). At the same time, the eigenvectors were reproduced rather well (Fig. S8). Moreover, the problematic eigenmodes mostly involved the very flexible intra-base pair coordinate buckle, presumably less relevant in many applications.

To deal with the problem, we decided, after some experiments, to introduce a cutoff on the eigenvalues. We factorized the assembled stiffness matrix *K* as *K* = *PDP^T^*, where *D* is the diagonal matrix containing the eigenvalues and *P* is an orthogonal matrix whose columns are the unit eigenvectors (polar decomposition). Now the eigenvalues in *D* smaller than the cutoff *λ_c_* were replaced by *λ_c_*, obtaining a new diagonal matrix *D̃*, and the new stiffness matrix was computed as *K̃* = *PD̃P^T^*. An analogous method was proposed to construct a nearest positive semidefinite matrix to a given symmetric matrix, where negative eigenvalues were replaced by zeros (151). Thus, we believe that this method yields a stiffness matrix with the desired properties by introducing a minimal perturbation to the original one.

To optimize the cutoff, we cannot use the hexameric set107, since the matrix blocks were deduced from there and, if they are assembled back, they simply give the same banded matrices they were cut out from (save possibly of the duplicate hexamers, Methods). We cannot use the pentameric set52 either, since this is the validation set. Thus, we made use of the set of DNA and RNA oligomers containing all tetramers (set14), of which the DNA version has been published (65) and the RNA version was simulated for this work. The target quantities were the global material constants (stretching and twisting stiffness, TS coupling, dynamic bending and twisting persistence lengths). For a given cutoff, we assembled the stiffness matrices for the set52 sequences, generated the corresponding structural ensembles by a Monte Carlo method, then computed the material constants for this model data as well as for the actual MD data of set52. We repeated the procedure for various cutoffs and chose the one for which the Pearson correlation coefficients between the material constants computed in both ways were maximal. It has turned out that the correlations are weak functions of the cutoff (Figs S10, S11) and that *λ_c_* = 0.48 for DNA and *λ_c_* = 0.44 for RNA are good choices. With these cutoff values, the correlation coefficients against the validation set52 are high, 0.8-0.9 in most cases (Figs S12, S13). Furthermore, the mean relative error on the eigenvalues themselves are 4.1 % for DNA and 2.9 % for RNA. This indicates that the model stiffness matrices can reproduce both local and global flexibility of DNA and RNA duplexes rather well.

### Applications of the model

We have developed a model of sequence-dependent structure and harmonic deformability of DNA and RNA duplexes and parameterized it from all-atom, explicit-solvent MD simulation data of 107 DNA and 107 RNA oligomers containing all hexanucleotide sequences. Here we present some applications of the model to demonstrate its utility.

#### Signature triple minimum of minor groove profile at the Exd-Scr binding site

Transcription factors may recognize their target DNA sequences via specific, sequence-dependent shape or deformability features (shape readout (13)). Of these, the *Drosophila* Hox protein Sex combs reduced (Scr) has distinct DNA recognition properties when it binds as a heterodimer with its cofactor Extradenticle (Exd) (72). The crystal structure of the Exd-Scr complex bound to DNA (2R5Z (152)) suggests three locations of insertion from Exd and Scr arginine residues into the minor groove. The deep-learning based Deep DNAshape tool (22) predicted minor groove widths pre-formed (narrowed) at these three sites, while only two of them were identified by the earlier pentamer-based method (20).

Here we examined top ten of the highest-affinity sequences for the Exd-Scr complex with DNA reported in (72). Although the individual sequences give a somewhat noisy picture, their average clearly shows the signature triple minor groove minimum of the target DNA sequence (Fig. 6). Thus, our heptamer-based sliding window approach, parameterized from the MD data in a straightforward manner, can detect this subtle effect.

**Figure 5.**
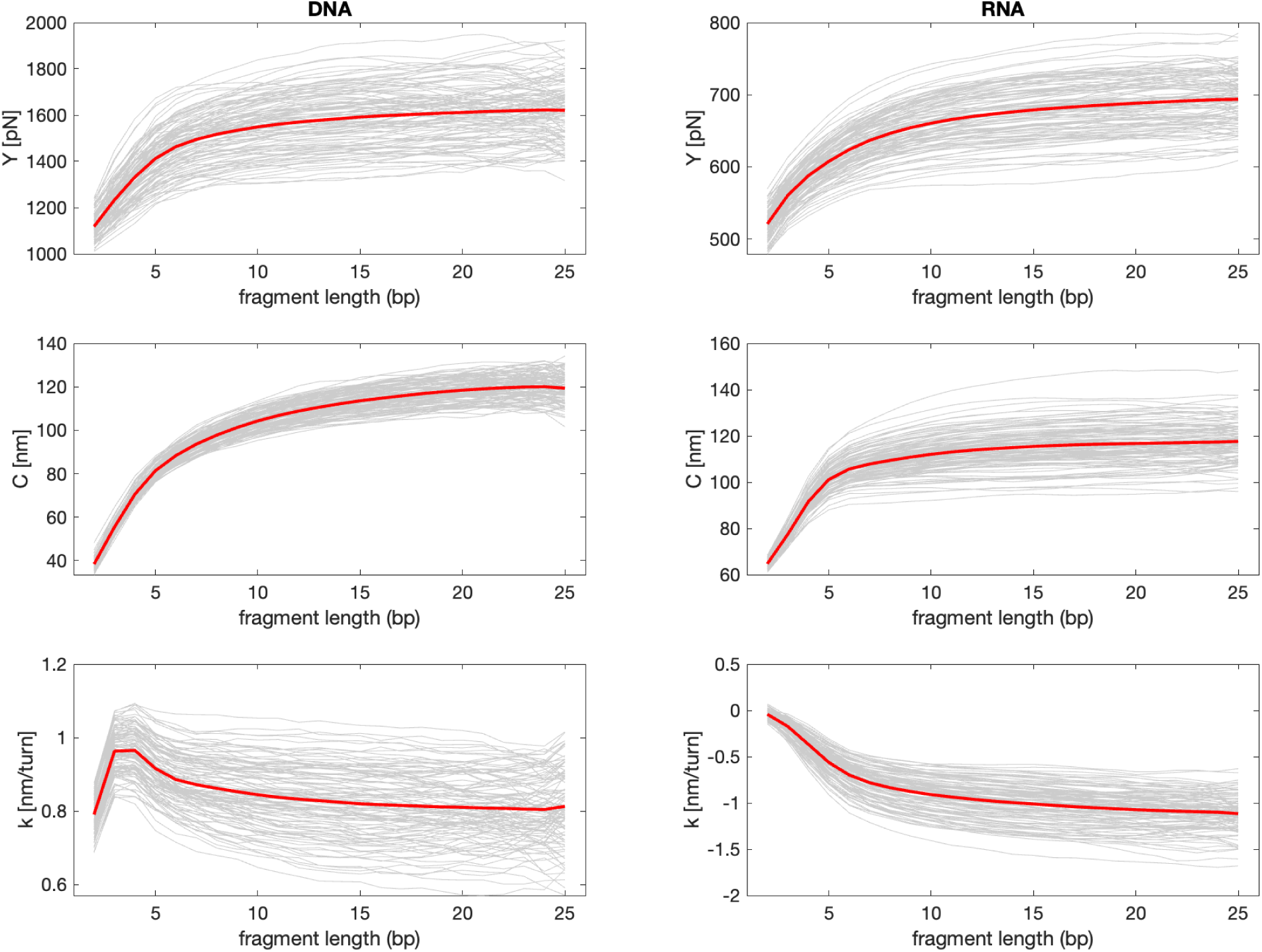
Length dependence of the elastic constants. The central 25 bp part of the simulated 107 DNA and 107 RNA oligomers containing all hexamers, and the fragments of this part, are examined, the values for fragments of the given length are averaged (grey lines). The red lines indicate sequence-averaged values (i.e. means over the data in grey). A universal, sequence-independent pattern emerges, with the stretch modulus and twist stiffness increasing with length and the twist-stretch coupling exhibiting a non-monotonic dependence. The 25 bp scale is about the minimal one to obtain converged values.

**Figure 6.**
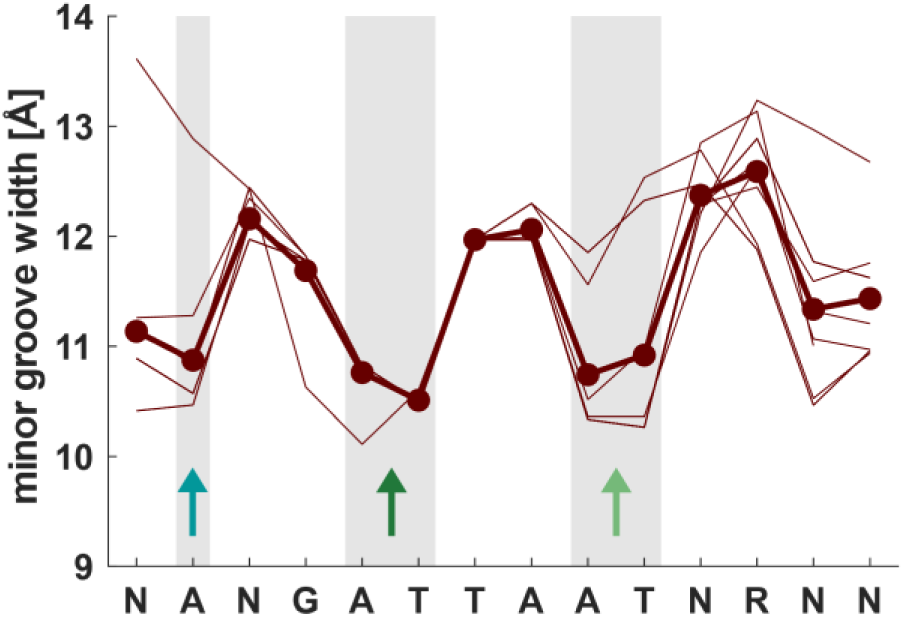
Predicted minor groove profiles for the Exd-Scr binding site. The profiles for top ten highest-affinity sequences (72) (thin lines) are somewhat noisy, but their average (in bold) exhibits the signature triple minimum associated with arginine residues from Exd (blue) and Scr (green and dark green) protruding into the groove (22). This prediction of our heptameric sliding window model is consistent with the one of Deep DNAshape (22) while an earlier, purely pentamer-based model gave just two minima (20).

#### Nucleosome destabilization by DNA polyA sequences

Nucleosomes are destabilized by polyA DNA tracts, with profound implications on genome function and chromatin organization (79,153,154). Using a non-local harmonic stiffness model parameterized from all-atom MD simulations of DNA sequences with short A-tracts, our group reported that the deformation energy of A-tracts in the nucleosome is position-dependent and is higher on average than that of a control sequence without A-tracts (80). However, the nucleosome destabilization is known to be polyA length-dependent and is more pronounced for longer tracts, such as A_34_ (155).

Here we designed a 401 bp random sequence (control, without A-tracts) and then mutated its central part to A_n_ (*n* = 21, 41, 61). The sequence-dependent structure and deformability was estimated using the model described above (hexameric/heptameric sliding window for the structure, stiffness matrix assembly from hexanucleotide blocks, with the cutoff applied). We threaded the sequences through a nucleosome structure (1kx5 (156)) and computed the deformation energy as in Eq. 1. We only considered the step coordinates roll, twist and slide, since these are highly conserved among nucleosome structures (157). All the other coordinates were left free to adopt energetically optimal values (partially relaxed model (80)). The mechanical destabilization by the A_n_ sequences is clearly visible (Fig. S14). Averaging the energetic cost over the positions where the whole A_n_ tract is within the nucleosome, we find the energy difference Δ*E* proportional to the tract length and close to 1 kcal/mol per base pair. Thus, our model correctly predicts the nucleosome destabilization by the polyA DNA sequences, and provides an estimate of the associated energetic cost.

#### Contrasting bending mechanisms of DNA A-tracts and RNA AU-tracts

It has long been known that A-tracts induce a bend to the DNA double helix towards the minor groove at the tract centre, although the exact magnitude of the bend and its dependence on the ionic environment are still debated (6,79,158–161). Structural hallmarks of the DNA A-tracts are a narrow minor groove and a high negative propeller twist. A recent study reported global bending also for the RNA double helix, this time caused by the (AU)_n_ sequences (AU-tracts): atomic force microscopy imaging showed that phased AU-tracts induce a macroscopic curvature, while all-atom MD simulations identified a localized compression of the dsRNA major groove and a large negative propeller twist at the tracts (12). These data suggest that there may be similarities as well as differences between the two bending phenomena.

Here we constructed a 28 bp random G/C sequence in its DNA and RNA form, and mutated its centre to A_8_ for DNA and (AU)_4_ for RNA, respectively. The key predictions are in Fig. 7. The coloured bands are values within one standard deviation from the mean over all hexameric (for the roll) or heptameric sequences. This enables one to quantify the notion of extreme values, meaning values outside this interval. The A-tract is predicted essentially straight (small roll) with higher roll values at its borders or in the non-A-tract sequence, in line with many earlier reports (summarized in (79)). By contrast, the AU-tract exhibits very high positive roll in UA steps accompanied by small, but still positive roll in AU steps. The DNA A-tract minor groove is narrow, as well known (79), while our model also predicts an expanded minor groove in the RNA AU-tract. The model reproduces a compressed major groove of the AU-tract (Fig. S15), and a high negative propeller in both tracts, as reported (12). However, here we also examine the propeller stiffness and find a striking difference: the A-tract propeller is exceptionally stiff (high force constant, Fig. 7D), while the AU-tract propeller is very flexible (low force constant). Finally, we notice the gradual build-up of the A-tract structure in the 5’-3’ direction saturated only at the 4^th^ adenine, the minimal A-tract length (79) (Fig. 4B-D), a phenomenon not present in the RNA AU-tract.

**Figure 7.**
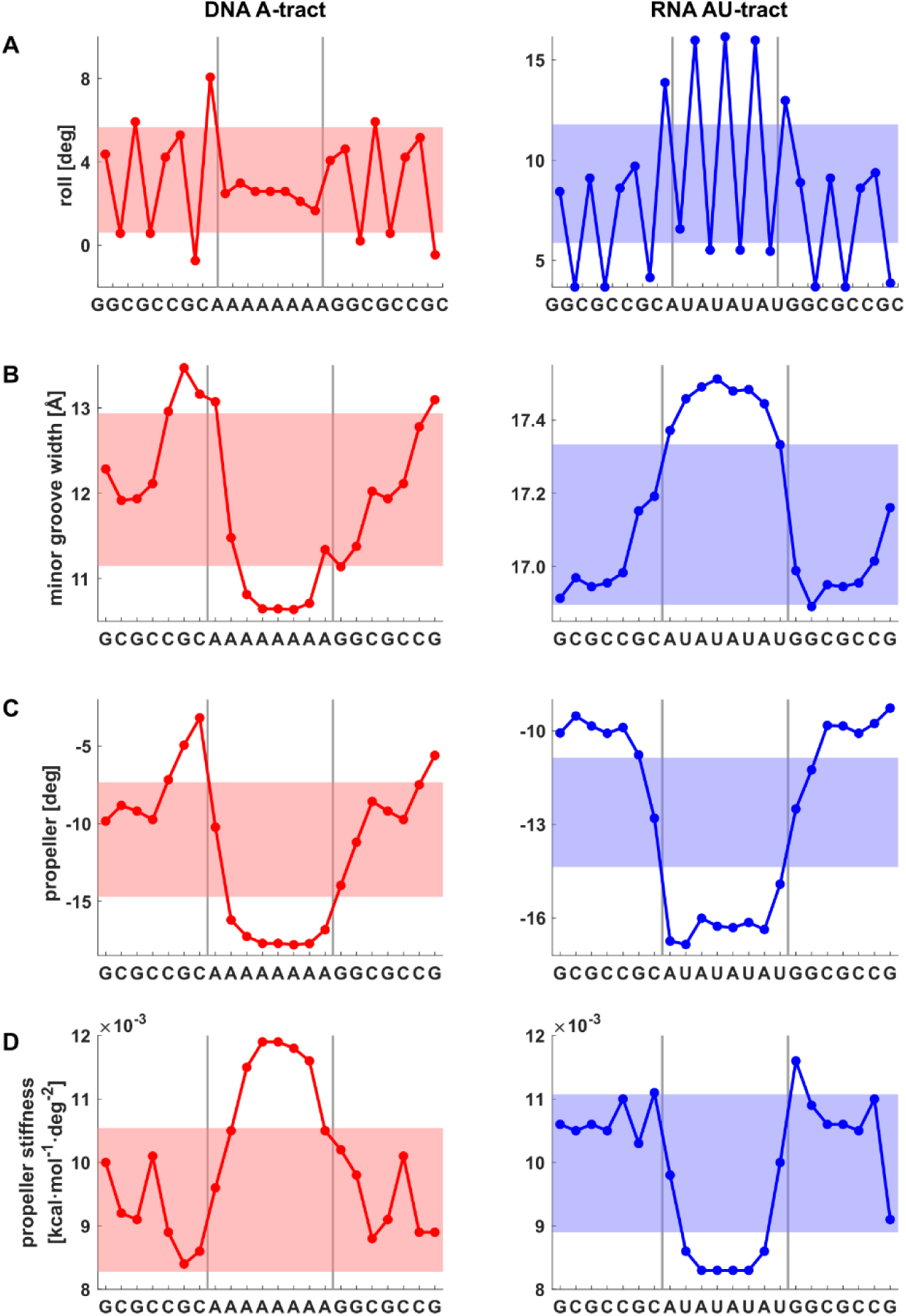
Contrasting profiles of sequence-dependent shape and stiffness parameters for DNA A-tract and RNA AU-tract sequences. The coloured stripes indicate values within one standard deviation from the mean over all hexameric (roll) or heptameric sequences. The values suggest the RNA AU-tract to be bent towards the major groove, non-cooperative, and flexible in propeller, in stark contrast to the DNA A-tract.

Taken together, these data suggest contrasting bending mechanisms for DNA A-tracts and RNA AU-tracts. The A-tract appears as a straight, cooperative, rigid unit, exhibiting narrow minor groove and a high negative, stiff propeller twist, the helix bending towards the minor groove being localized mostly outside the tract or at the junctions, in line with earlier models (79). By contrast, we find the AU-tract flexible in propeller, non-cooperative and itself highly bent, this time towards the major groove it its centre. The bending is due to the high positive roll in UA steps, further supported by smaller but still positive roll in AU steps and accompanied by the expanded minor groove. Of note, an analogous bending mechanism of a DNA AT-tract is not possible due to a much smaller positive roll in TA steps, further neutralized by the negative roll in AT steps (Fig. S16).

### Structural dynamics

The amount of the simulated MD data (107 DNA oligomers, each simulated for 2 µs, and 107 RNA oligomers, 1 µs each) enabled us to detect rare events, such as base-pair opening or alternative (non-canonical) structures. We focused on the central 25 bp parts containing all hexanucleotide sequences, the GCGC caps were omitted.

The base pairs occasionally open for a short time, then close again. We consider a pair open if at least one of the Watson-Crick hydrogen bonds is broken, by which we mean that the distance between the heavy atoms is greater than 4 Å, as proposed in (46). A proper representation of hydrogen bonding is critical especially for folding non-canonical RNA structures and is still a focus of intense research (104,162,163).

In the duplexes studied here, the population of broken A-T and G-C pairs is very small in all cases: 0.15 and 0.10 % respectively for DNA, 0.38 and 0.63 % for RNA. Thus, the RNA pairs appear less stable in MD than the DNA ones, and G-C less so than A-T. Notice that we do not monitor a complete flip-out of a base, but a smaller perturbation, in the spirit of imino proton exchange measurements where the opening angle as small as ± 30° is sufficient for the exchange to take place (164).

Next, we examined the open pair lifetimes. Rather than blindly averaging the timespans of all opening events, we adopt (and simplify) a method proposed in (165). The opening times are considered as independent, identically distributed random variables. This is in line with the experimental finding that base pairs open one at a time, but neglects the dependence on base-pair identity (A-T vs. G-C) and on sequence context (166). Under this assumption, we construct the survival function *S*(*t*) which represents the probability that the pair remains open longer than time *t*. If *S*(*t*) = exp(−*λt*), then the mean opening time is *τ_open_* = − 1⁄*λ* (165). We computed *S*(*t*) over all opening events, separately for DNA and RNA duplexes, and plotted the logarithm of *S*(*t*). For short times, ln *S*(*t*) is visibly non-linear, indicating brief transient escapes, and it is not well defined for long times (Fig. S17). In between however, there is a time interval where it is close to linear. Rather than computing the time derivative as proposed in (165), we just fitted a straight line there, whose slope *λ* then defines the mean opening time. We found *τ*_*open*_ of 2.3 ns for DNA and 5.3 ns for RNA. Thus, our MD data predict open pair lifetimes in DNA and RNA duplexes to be in the nanosecond range, and longer for RNA, in agreement with imino proton exchange measurements (166).

Finally, we scanned the time series of the intra-base pair and step coordinates to find time intervals of anomalous values, indicative of a non-canonical structure. In all the DNA MD data, we found only four such structures with lifetimes longer than 10 ns (Fig. S18). One of them showed inter-strand stacking of the pairs, three others were ladder-like structures associated with long-living concerted flips of the backbone torsions *α*/*γ* from *g+/t* to *g-/g+*, accompanied by the shift of *β* from *t* towards lower *g-* (around 240°).

No such structures were observed in the RNA duplexes. Instead, a spectacular flip of a sugar pucker into the B-DNA domain, lasting for 250 ns, was seen, associated with a flip of the torsion angle *κ* involving the 2’-OH group (H2’-C2’-O2’-HO2’) from wildly fluctuating around 50° to locked around 130° (Fig. 8). Changes in the propeller of the pair in question, as well as twists in the steps involved, were also observed (Fig. S19). Several other, shorter flips (< 50 ns) were detected. This type of RNA sugar pucker *κ* flip was recently identified in the *E. coli* sarcin-ricin loop using cryo neutron crystallography (167), but the residues involved were all extra-helical. Our MD data suggest that such flips may take place within the RNA duplex as well.

**Figure 8.**
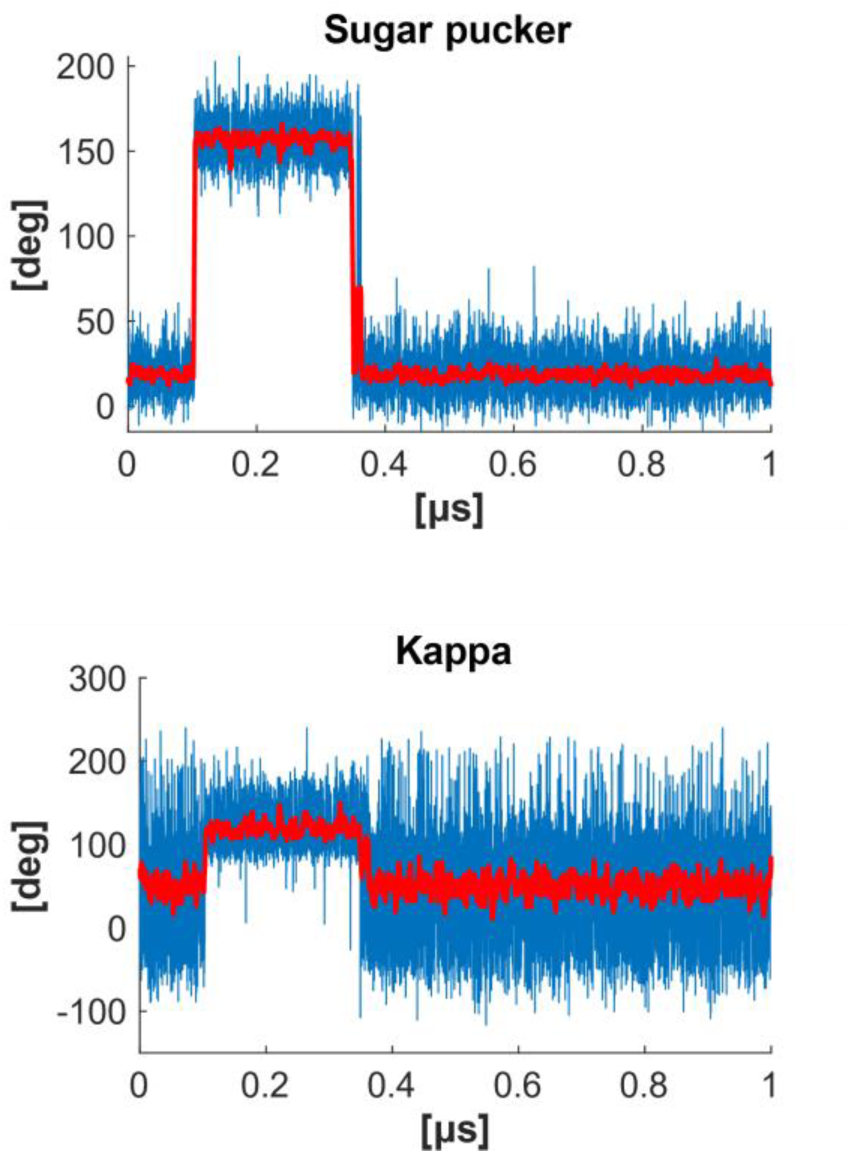
A long-living (250 ns) flip of the RNA sugar pucker into the B domain (RNA hexameric sequence set, seq. 79, U in the uauUguu context), accompanied by a flip in the torsion angle *κ* involving the 2’-OH group. Data averaged over a sliding window of 3 ns are shown in red. Several other, shorter (< 50 ns) flips of this kind were also detected in our MD simulations.

## Conclusion

It has long been recognized that the structure and deformability of the DNA double helix is modulated by its base sequence, with fundamental ramifications for DNA biology and nanostructure design. Despite intense research, this dependence has remained incompletely understood. Sequence-specific variations of shape and stiffness have been emerging also for double-stranded RNA, a prominent structural motif. In this work we performed a set of atomic-resolution, explicit-solvent MD simulations of double-stranded DNA and RNA oligomers containing all the 2080 unique hexanucleotide sequences, and analysed them in terms of sequence-specific shape and harmonic deformability. We identified two scales of effective base-base interactions within the DNA and RNA double helices and exposed fundamental differences in the rules governing DNA and RNA sequence-dependent structure and stiffness. We constructed a model to predict DNA and RNA shape and harmonic stiffness for arbitrary sequence, validated it on a dataset of MD-simulated oligomers involving all DNA and RNA pentameric sequences, and demonstrated its utility in various applications.

We then moved to the global level where the duplexes are described by a small number of material constants. The large, well-balanced set of oligomers enabled us to study the sequence dependence of the stretch modulus, twist rigidity, twist-stretch (TS) coupling, as well as the bending and twisting persistence lengths. The sequence-specific variability in these constants was around ±20 % (50 % for the TS coupling). Universal length dependencies of the constants were identified, while the static bending and twisting disorder was found to be very small.

Overall, the present work provides a comprehensive description of DNA and RNA duplexes at the hexanucleotide scale with minimum additional assumptions, proposes and validates a straightforward model to predict shape and harmonic stiffness for an arbitrary sequence, and probes sequence and length dependence at the global scale. The model parameters are made freely available, forming a baseline of further research and allowing for a broad range of applications in molecular biology, biophysics, and nucleic acid nanostructure design.

### Data availability

The model parameters, heatmap visualizations, as well as the in-house Python script to generate a minimal sequence including all *k*-mers, are available at Zenodo, https://zenodo.org/records/14936465.

## Supporting information

List of simulated oligomers containing all hexanucleotides

## Supplementary data

Supplementary data are available at NAR Online.

## Acknowledgements

The authors would like to thank Remo Rohs for useful discussions, and Tomáš Dršata for his participation in the initial stage of the project.

