## Supporting information for "Sequence-dependent shape and stiffness of DNA and RNA double helices: hexanucleotide scale and beyond"

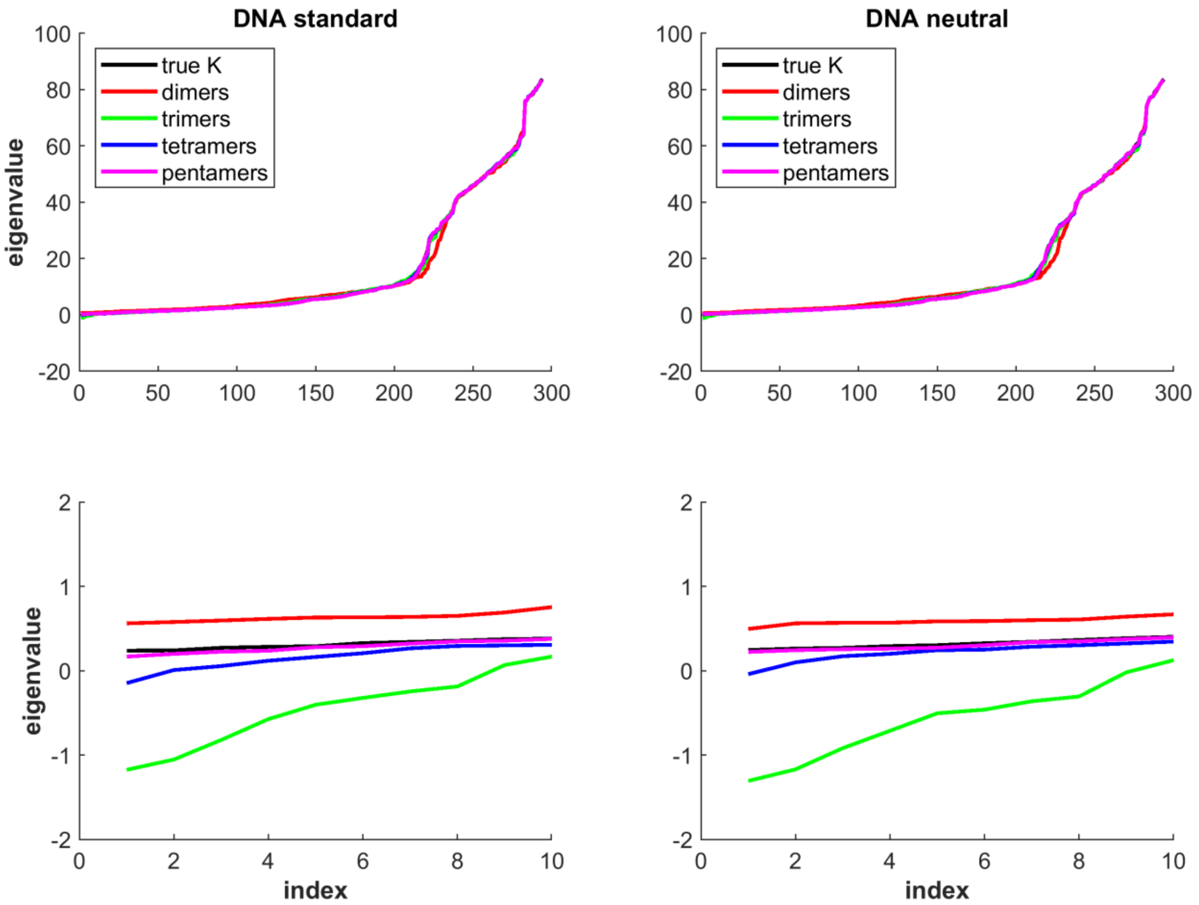


**Figure S1.** Eigenvalues of the (non-dimensionalized) stiffness matrix for the s0 sequence in its DNA form, with various base-base interaction ranges imposed (see main text).

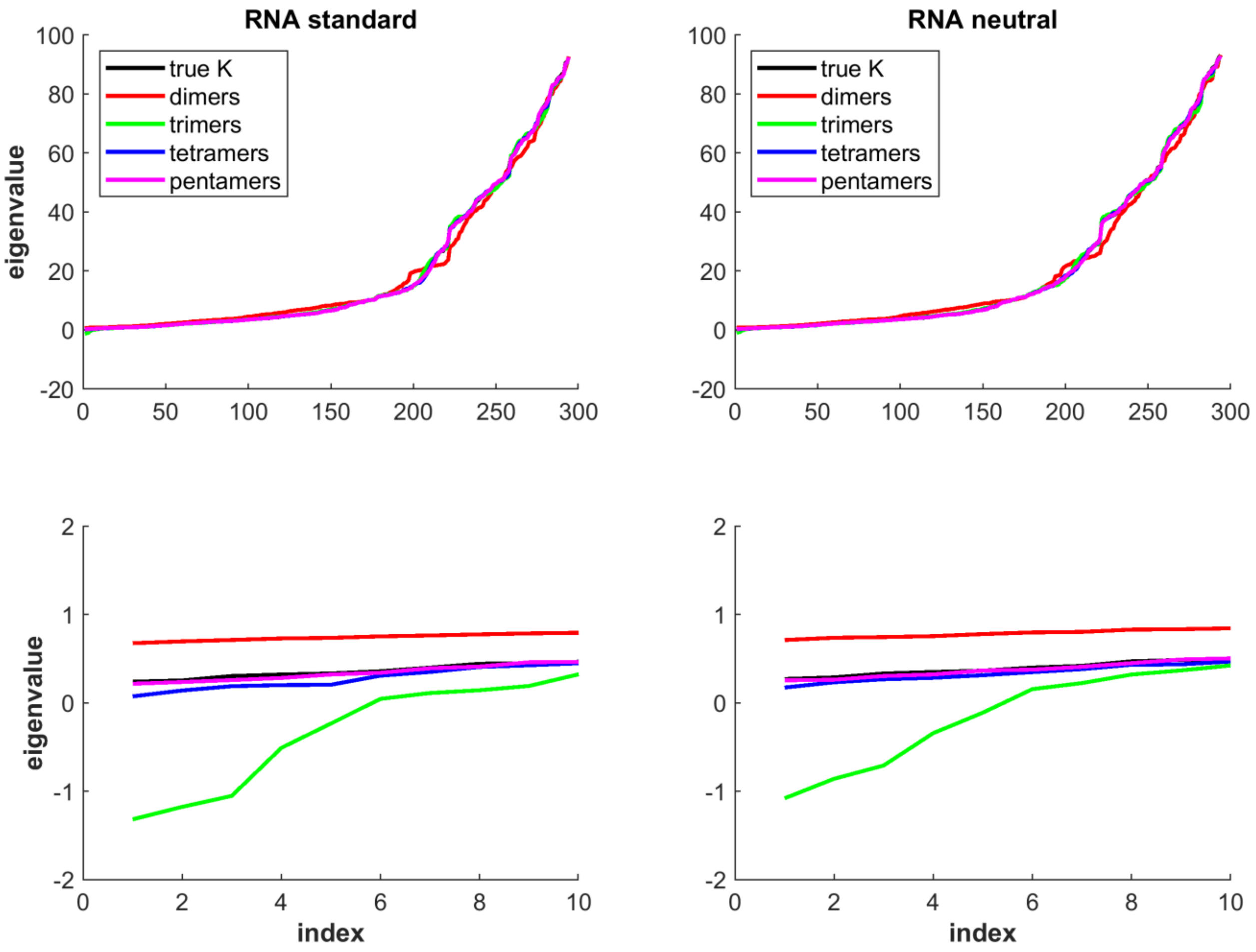


**Figure S2.** Eigenvalues of the (non-dimensionalized) stiffness matrix for the s0 sequence in its RNA form, with various base-base interaction ranges imposed (see main text).

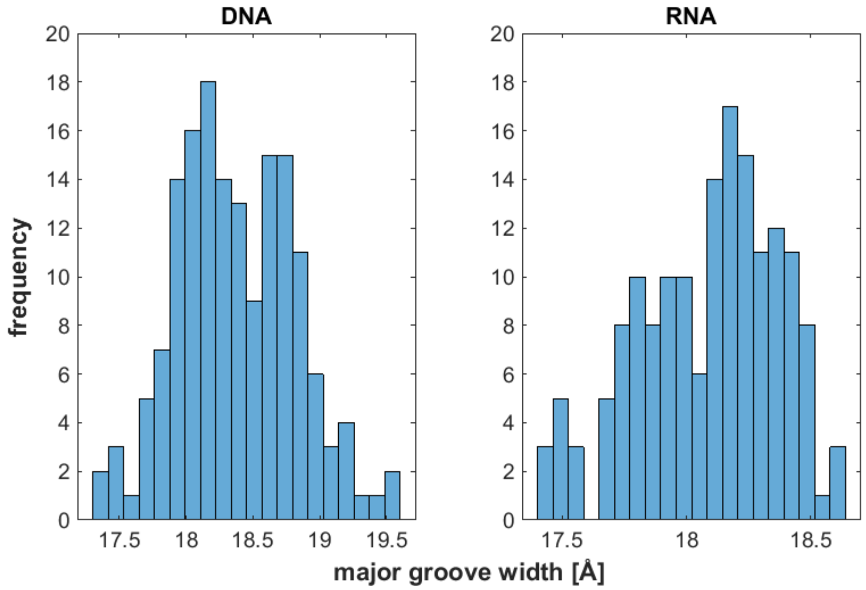


**Figure S3.** Histograms of major groove width for all the hexanucleotide sequences.

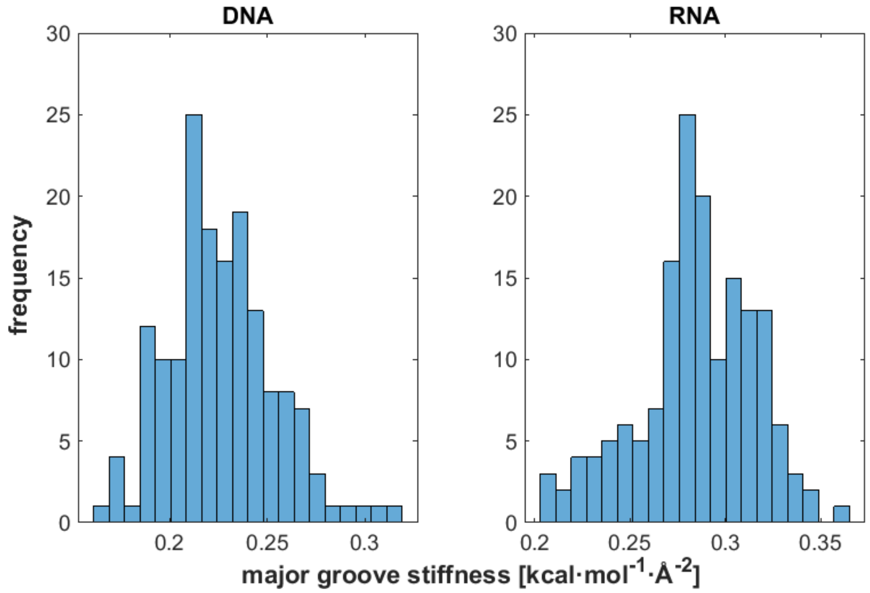


**Figure S4.** Histograms of major groove stiffness for all the hexanucleotide sequences.

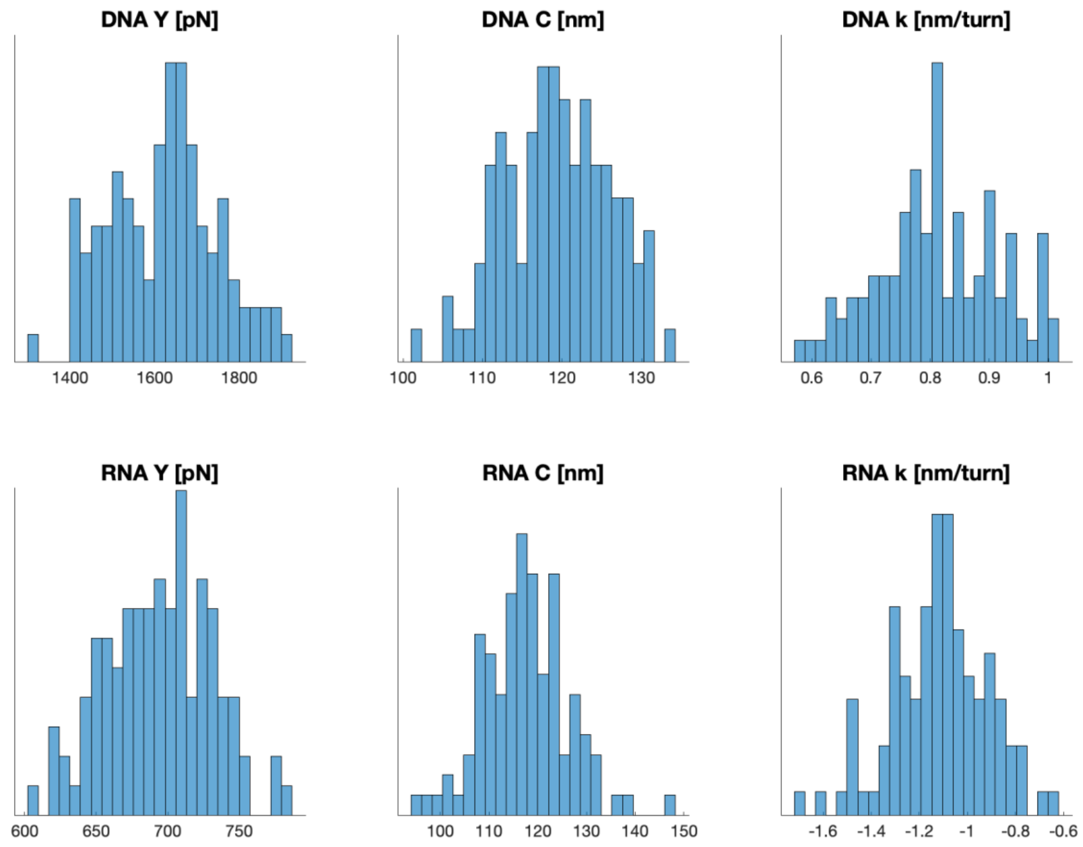


**Figure S5.** Histograms of the stretch modulus, twist rigidity, and twist-stretch coupling computed for the 107 DNA and 107 RNA oligomers containing all hexanucleotide sequences.

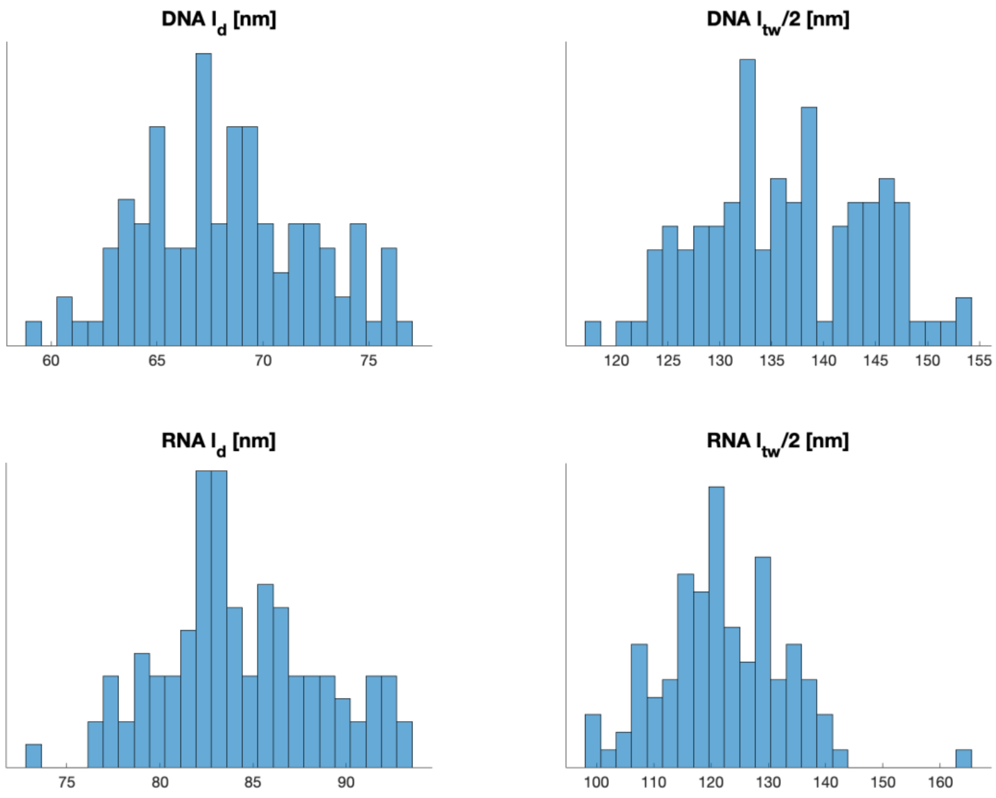


**Figure S6.** Histograms of bending and half twisting persistence lengths computed for the 107 DNA and 107 RNA oligomers containing all hexanucleotide sequences.

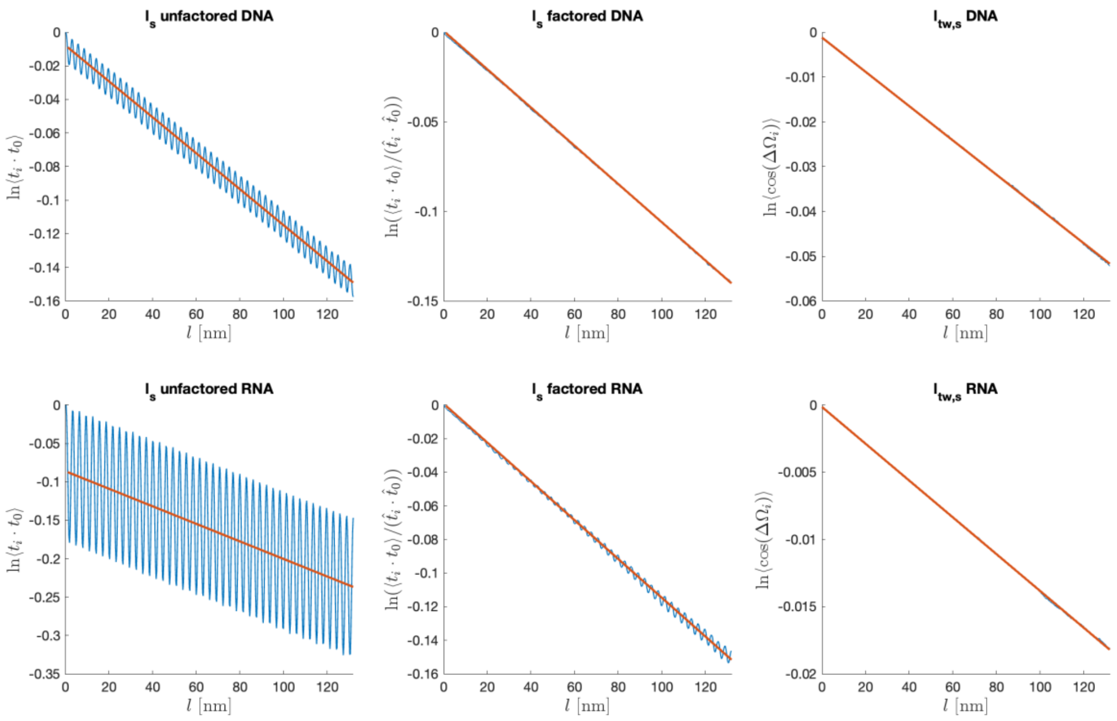


**Figure S7.** Semilog plots to deduce static bending and twisting persistence lengths.

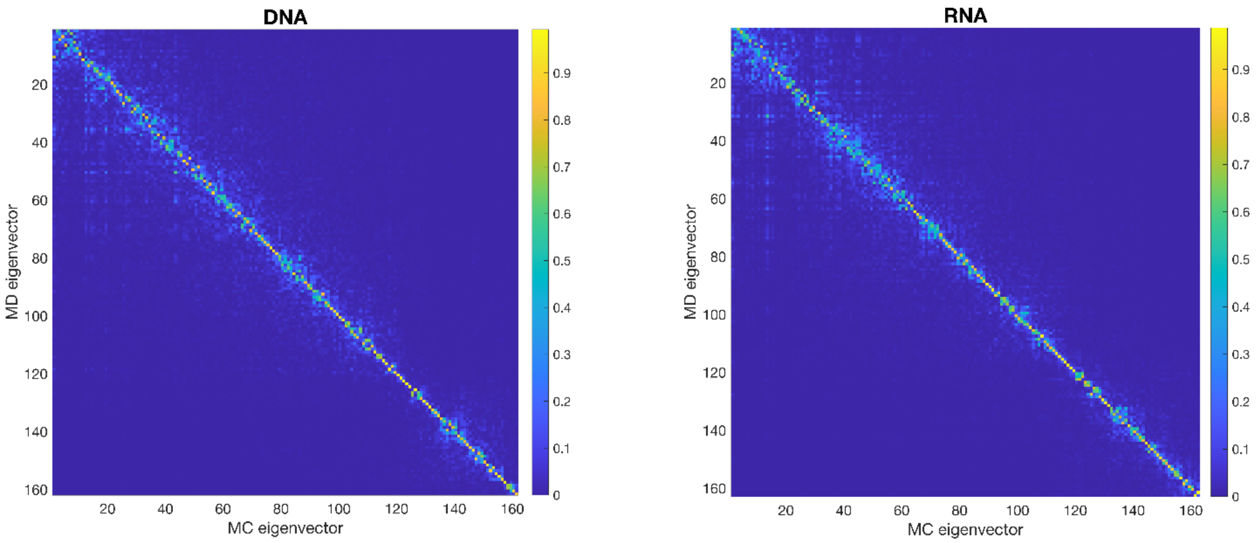


**Figure S8.** Absolute values of dot products between eigenvectors of a stiffness matrix assembled from hexameric blocks and those from the stiffness matrix inferred directly from MD data of the same sequence in the validation set52. Data for sequence 37 of set52 are shown as a typical example.

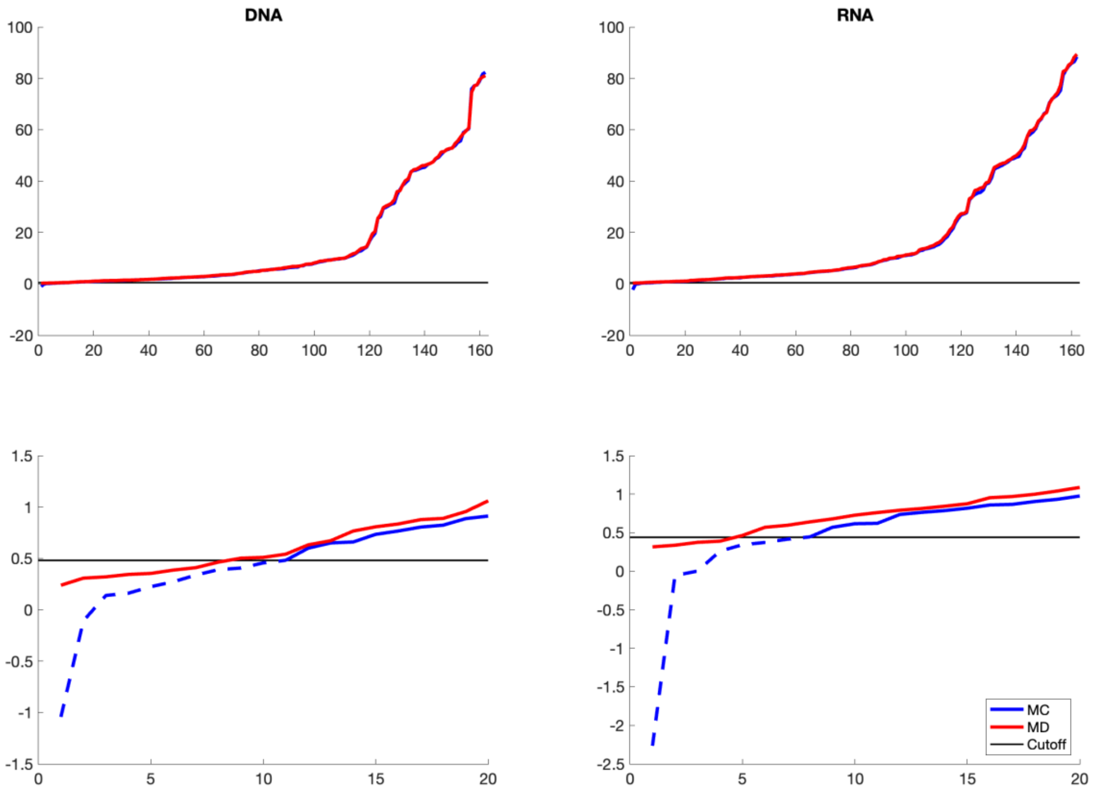


**Figure S9.** Eigenvalues of the MD-based stiffness matrix (seq. 37 of set52, red) and from its approximation by assembled hexameric blocks (blue). The smallest eigenvalues of the approximate matrix are close to zero and even negative and are replaced by the cutoff value (grey).

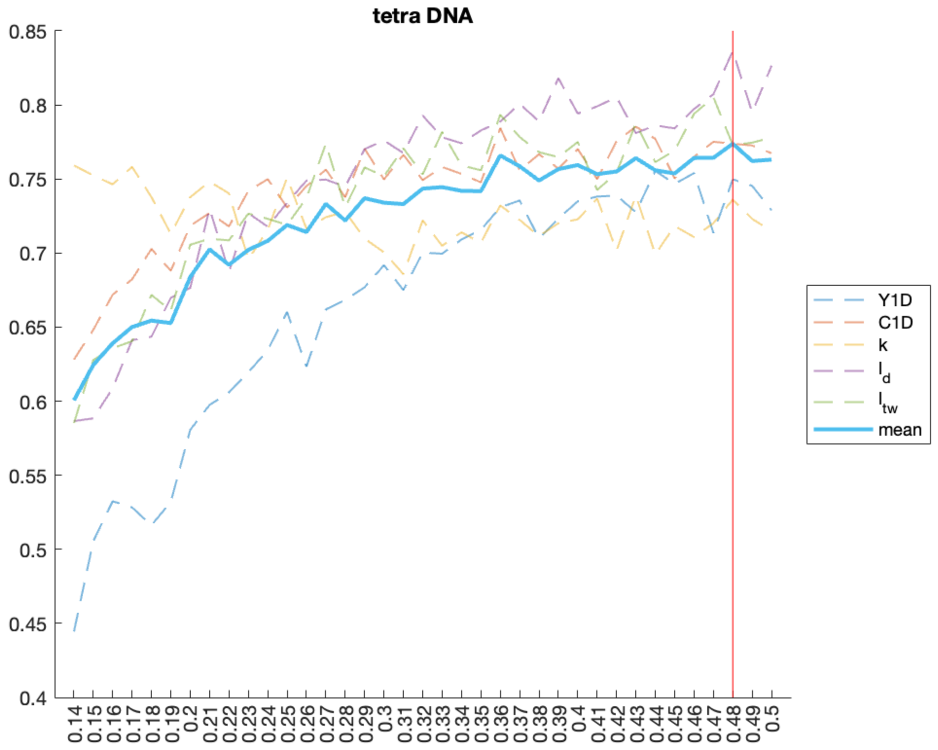


**Figure S10.** Pearson correlation coefficients between material constants deduced from MD set14 (containing all tetramers) and those from Monte Carlo generated structural ensemble using the approximate stiffness matrices (assembled from hexameric blocks) and the given cutoff (indicated on the *x* axis). The optimal DNA cutoff value was chosen to be 0.48 (red vertical line).

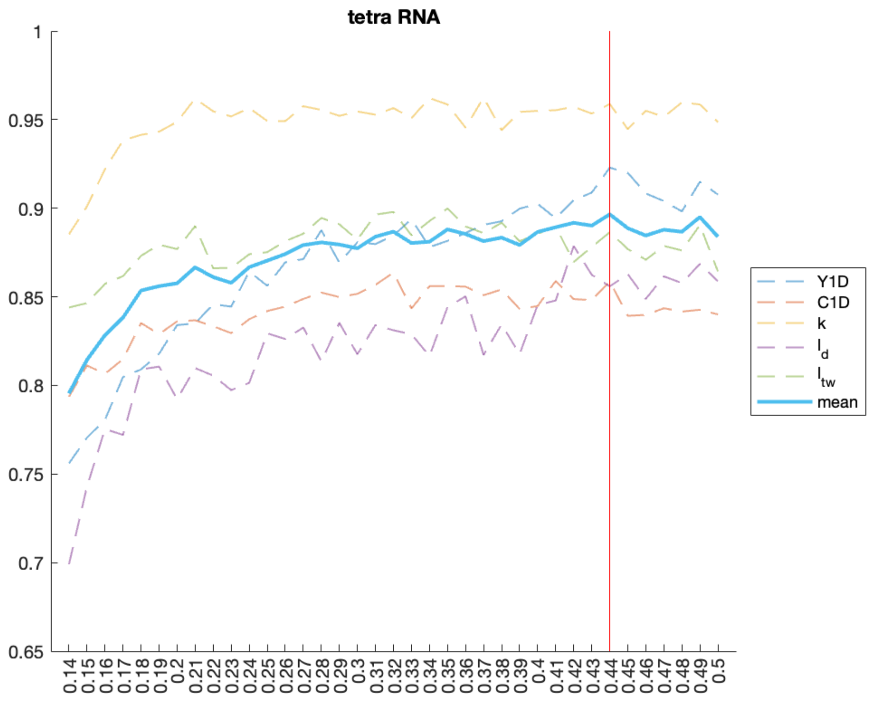


**Figure S11.** Pearson correlation coefficients between material constants deduced from MD set14 (containing all tetramers) and those from Monte Carlo generated structural ensemble using the approximate stiffness matrices (assembled from hexameric blocks) and the given cutoff (indicated on the *x* axis). The optimal RNA cutoff value was chosen to be 0.44 (red vertical line).

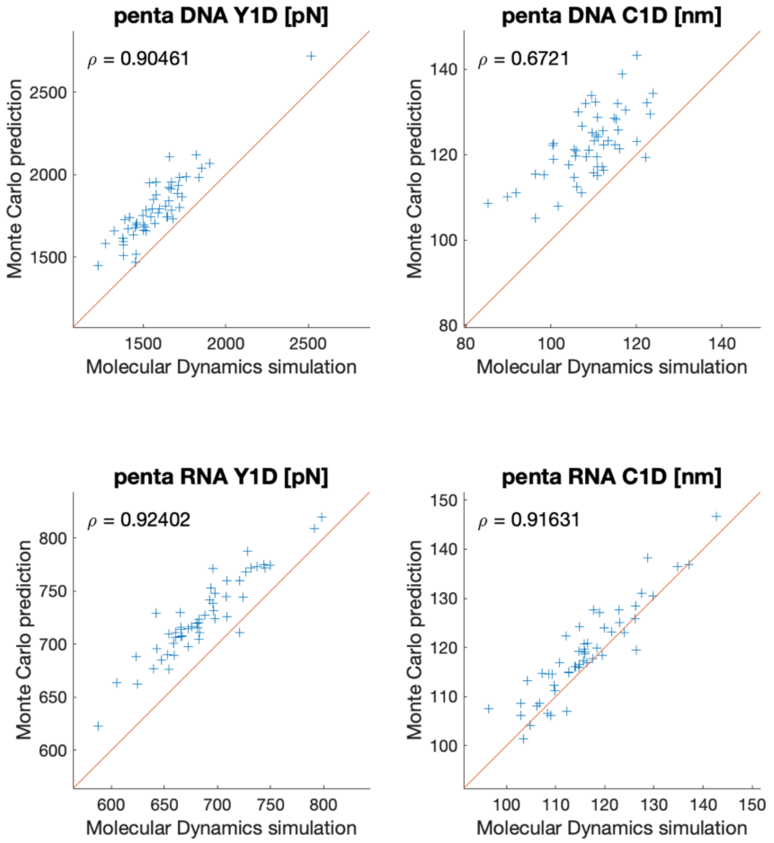


**Figure S12.** Correlations between elastic constants for the set52 sequences from MD and from MC simulations using the assembled stiffness matrices and the optimal cutoff.

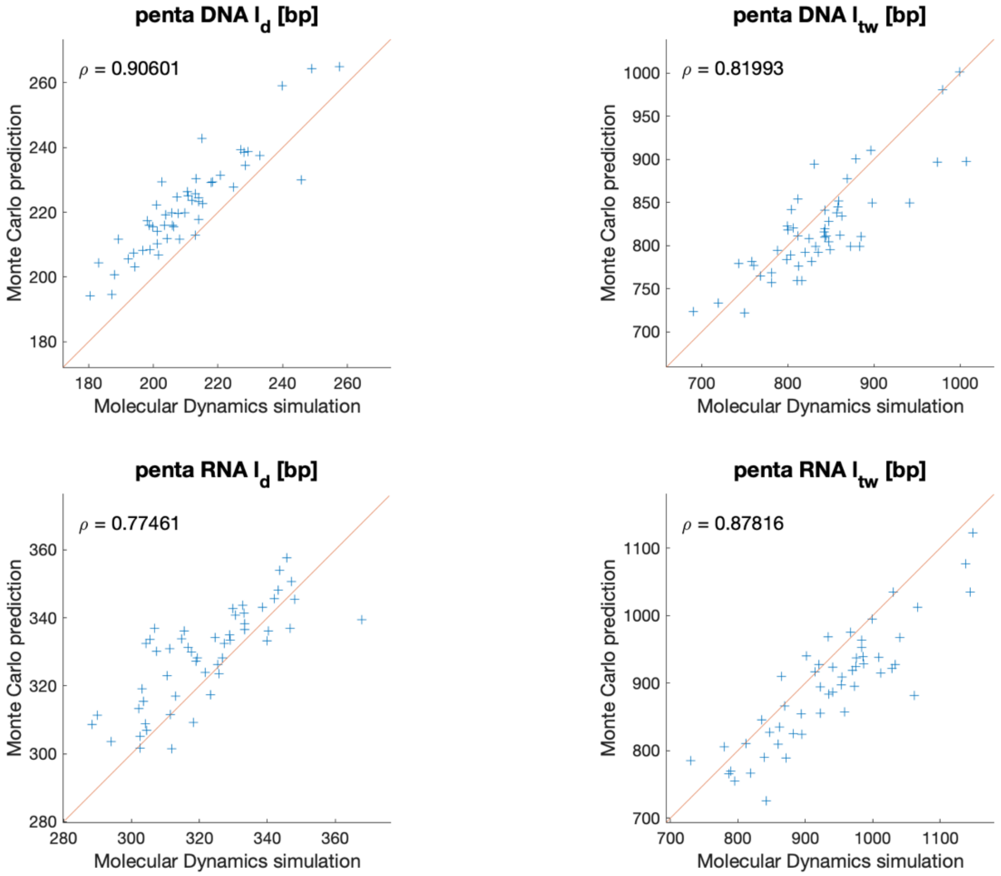


**Figure S13.** Correlations between persistence lengths for the set52 sequences from MD and from MC simulations using the assembled stiffness matrices and the optimal cutoff.

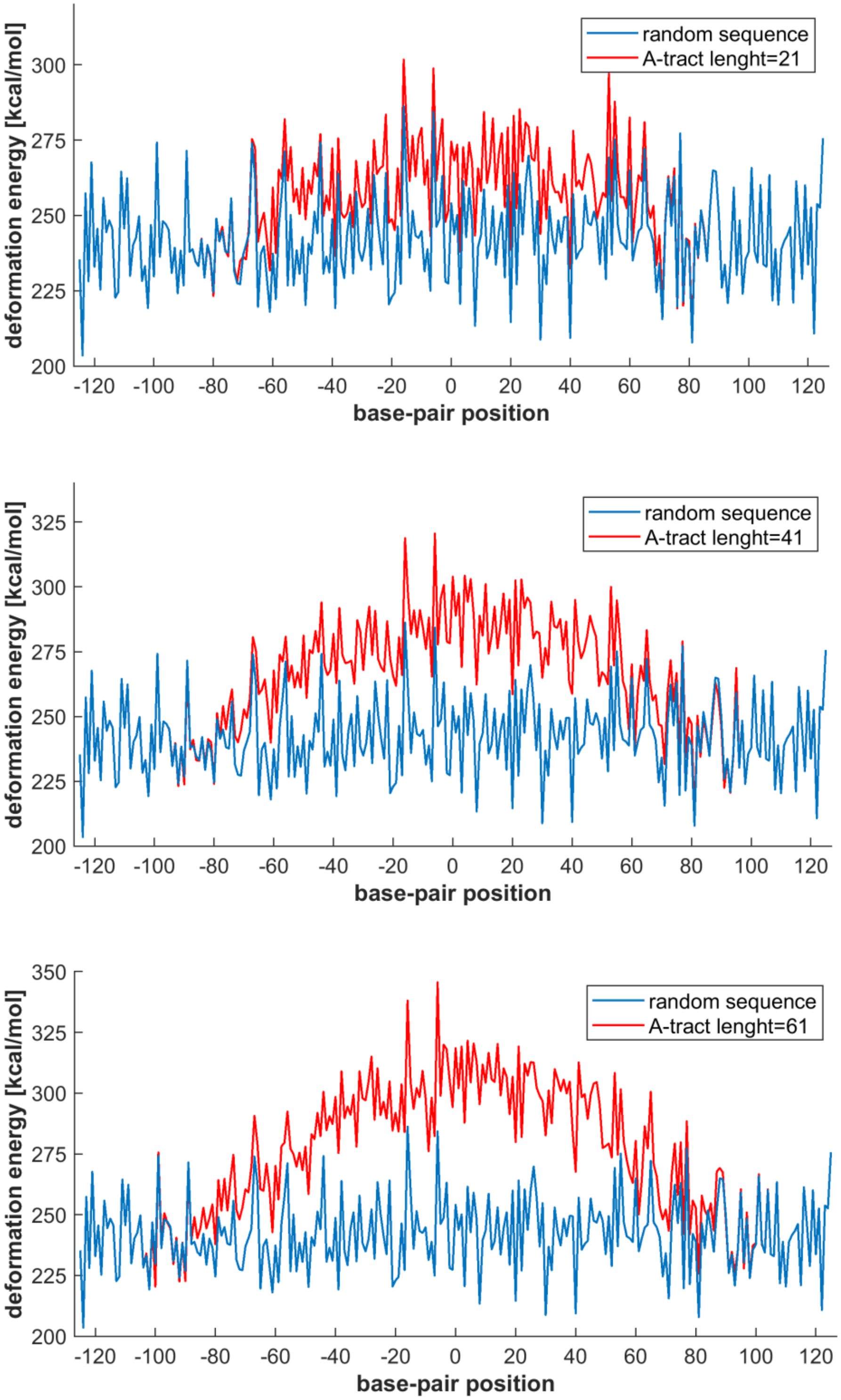


**Figure S14.** Deformation energy of threading a random sequence (blue), and the same sequence where the central part was mutated to polyA (red) through the 1kx5 nucleosome structure. The roll, twist and slide coordinates, highly conserved among nucleosome structures, were deformed, the remaining coordinates were relaxed to adopt energetically optimal values.

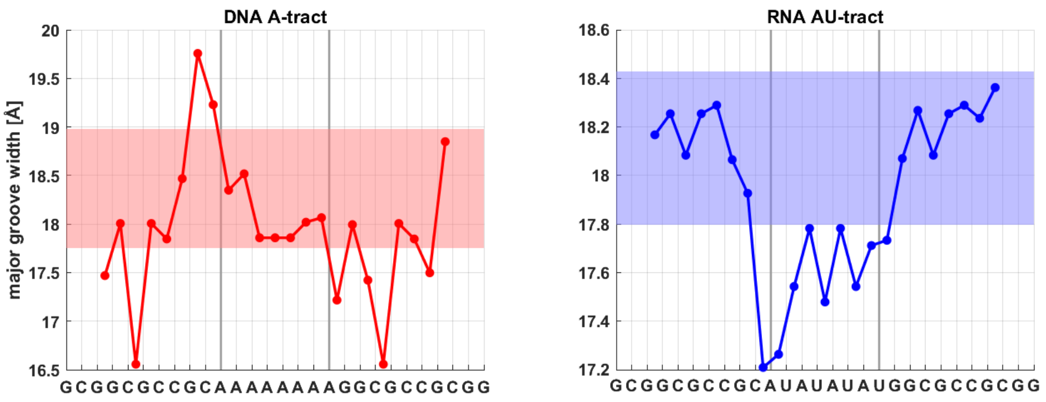


**Figure S15.** Major groove width profiles of a DNA A-tract and an RNA AU-tract as predicted by our model. The stripes indicate values within one standard deviation from the mean over all hexamers.

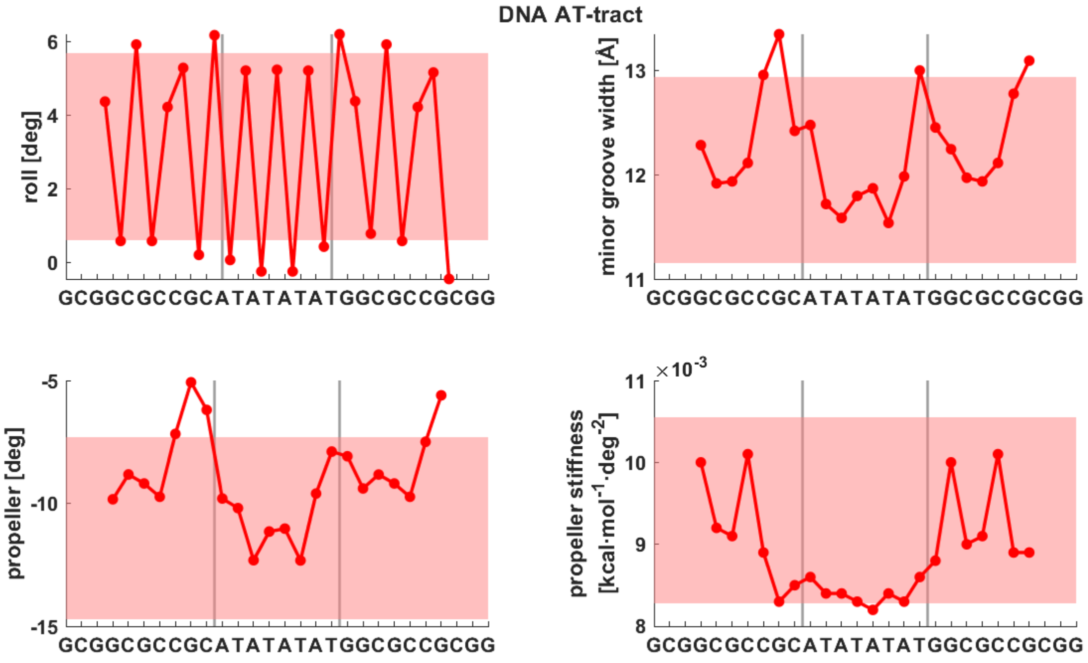


**Figure S16.** Coordinate profiles for a DNA AT-tract predicted by the model.

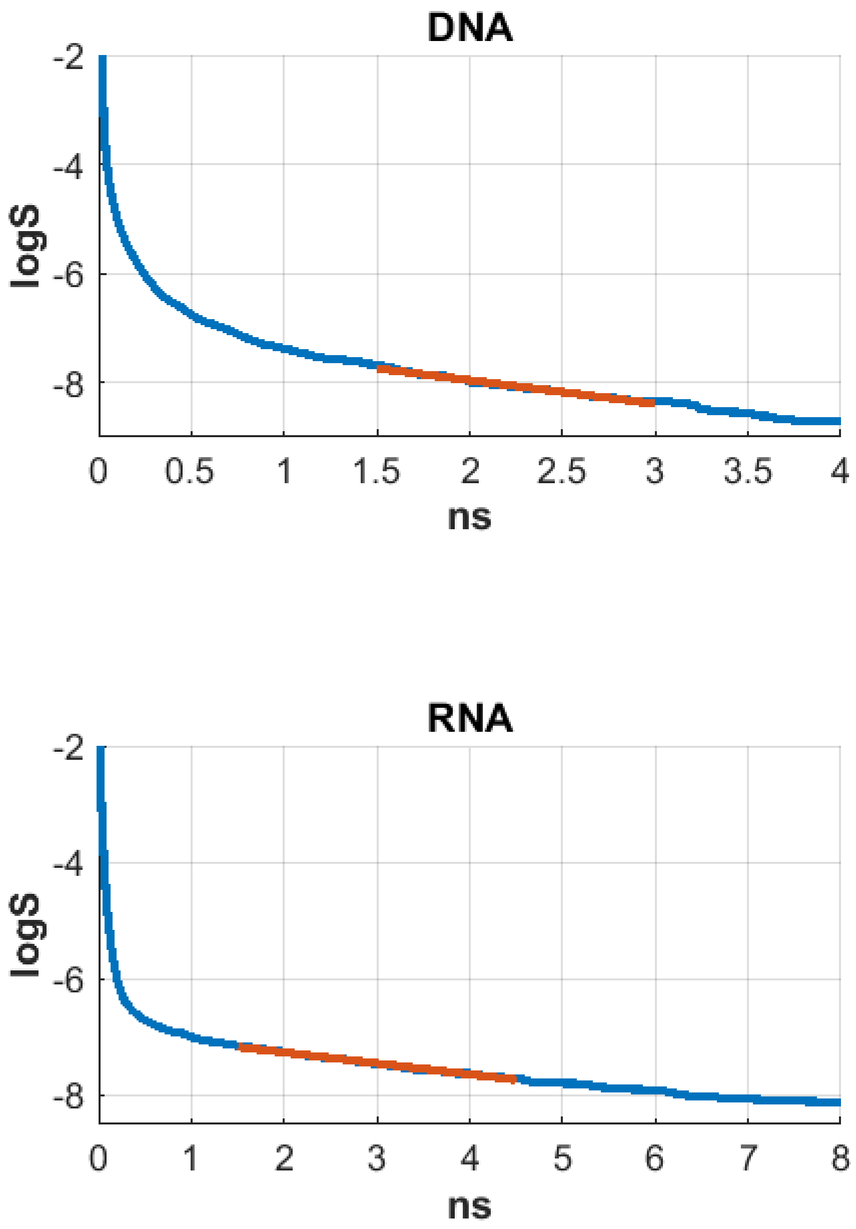


**Figure S17.** Logarithm of the base-pair opening survival function vs. time. The visibly linear part was fitted with a straight line (red) to obtain the mean opening times.

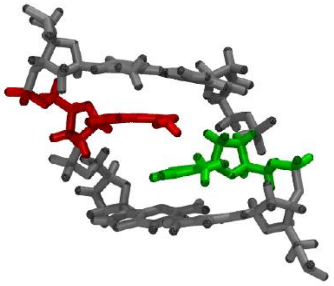

**Figure S18.** Two types of non-canonical structures observed in the 107 DNA simulation set, with lifetimes > 10 ns. An interstrand stack (left, one case) and a ladder-like structure associated with concerted flips of the backbone torsions (right, three cases).

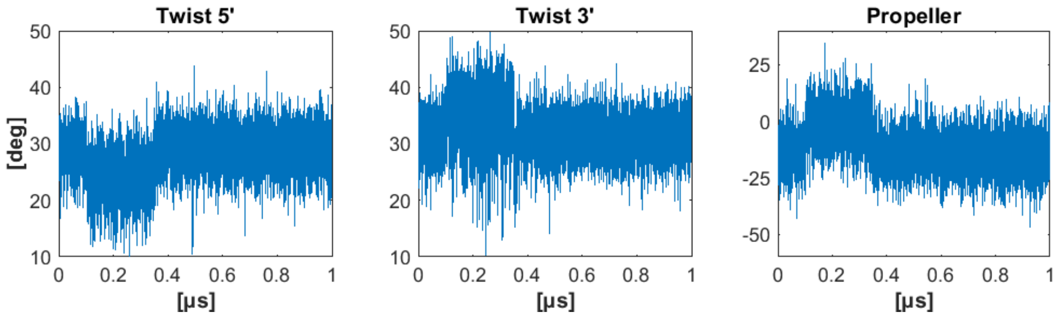


**Figure S19.** Time series of twist in the RNA steps surrounding the pair with the propeller flipped into the B domain, and the propeller of the pair itself. Significant changes are observed in the time interval of the flip (100 – 350 ns). Interestingly, the sum of the twists still attains the usual value, so that the RNA duplex as a whole is not over- or underwound during the flip.

**Table S1.** Errors on the equilibrium intra-base pair coordinates and minor groove widths predicted by the heptanucleotide model compared to the actual MD values from the set52 validation dataset.

| | **DNA** | **RNA** |
|---|---|---|
| Shear [Å] | 0.005 | 0.005 |
| Stretch [Å] | 0.002 | 0.002 |
| Stagger [Å] | 0.017 | 0.014 |
| Buckle [°] | 0.983 | 0.581 |
| Propeller [°] | 0.623 | 0.392 |
| Opening [°] | 0.108 | 0.080 |
| Minor g. width [Å] | 0.162 | 0.046 |

**Table S2.** Errors on the equilibrium inter-base pair and helical coordinates, and major groove widths, predicted by the hexanucleotide model compared to the actual MD values from the set52 validation dataset.

| | **DNA** | **RNA** |
|---|---|---|
| Shift [Å] | 0.042 | 0.016 |
| Slide [Å] | 0.038 | 0.017 |
| Rise [Å] | 0.025 | 0.017 |
| Tilt [°] | 0.186 | 0.154 |
| Roll [°] | 0.251 | 0.248 |
| Twist [°] | 0.457 | 0.141 |
| X-disp [Å] | 0.082 | 0.032 |
| Y-disp [Å] | 0.067 | 0.034 |
| h-Rise [Å] | 0.024 | 0.016 |
| Inclination [°] | 0.419 | 0.395 |
| Tip [°] | 0.315 | 0.280 |
| h-Twist [°] | 0.448 | 0.181 |
| Major g. width [Å] | 0.264 | 0.106 |

**Table S3.** Relative errors on 1D stiffness constants compared to the validation set s52.

| | DNA | RNA |
|---|---|---|
| Shear | 4.17 % | 1.99 % |
| Stretch | 3.70 % | 1.65 % |
| Stagger | 2.50 % | 1.56 % |
| Buckle | 1.36 % | 1.03 % |
| Propeller | 2.55 % | 1.58 % |
| Opening | 4.58 % | 1.85 % |
| Minor g. | 1.35 % | 0.27 % |

| | DNA | RNA |
|---|---|---|
| Shift | 0.72 % | 0.63 % |
| Slide | 0.95 % | 0.78 % |
| Rise | 1.20 % | 0.99 % |
| Tilt | 3.40 % | 1.42 % |
| Roll | 2.36 % | 1.41 % |
| Twist | 1.20 % | 0.91 % |
| Major g. | 1.44 % | 0.59 % |

**Table S4.** Simulated oligomers comprising all 512 unique DNA pentamers.

| DNA | 5'-------------------3' | DNA | 5'-------------------3' |
|---|---|---|---|
| seq1 | GCGCAAAATAGCCACTCTGCGC | seq27 | GCGCGGAAACGCTTTTTCGCGC |
| seq2 | GCGCCTCTCTTCATAGTTGCGC | seq28 | GCGCTTTCTTGTGCATACGCGC |
| seq3 | GCGCAGTTTTGCCAGAGGGCGC | seq29 | GCGCATACCAACGGTGCTGCGC |
| seq4 | GCGCGAGGCACTGTTTACGCGC | seq30 | GCGCTGCTCTATATCCCTGCGC |
| seq5 | GCGCTTACTAATCTCATTGCGC | seq31 | GCGCCCCTGGGCTGGAAGGCGC |
| seq6 | GCGCCATTATACAAATCAGCGC | seq32 | GCGCGAAGCTCCTGTGGTGCGC |
| seq7 | GCGCATCAATCGCATCTTGCGC | seq33 | GCGCTGGTGAACGTCTAGGCGC |
| seq8 | GCGCTCTTAGAAGGGCATGCGC | seq34 | GCGCCTAGCTGAAAGTCTGCGC |
| seq9 | GCGCGCATTCAAGCCTGAGCGC | seq35 | GCGCGTCTCCAAGGACGCGCGC |
| seq10 | GCGCCTGAGCGCGGTTGAGCGC | seq36 | GCGCACGCCTATCATGCTGCGC |
| seq11 | GCGCTTGAGTTCGTCAAAGCGC | seq37 | GCGCTGCTTACCTAACCAGCGC |
| seq12 | GCGCCAAACCCCAATTACGCGC | seq38 | GCGCACCAGTTGCGCCCGGCGC |
| seq13 | GCGCTTACACGCGAGGGGGCGC | seq39 | GCGCCCCGGAACCTTTAGGCGC |
| seq14 | GCGCGGGGCCGTGCGTAGGCGC | seq40 | GCGCTTAGCAAGTAGCGAGCGC |
| seq15 | GCGCGTAGTGTCGCCGGTGCGC | seq41 | GCGCGCGAATTTAATAAGGCGC |
| seq16 | GCGCCGGTACGACTGATCGCGC | seq42 | GCGCTAAGGCCATCCATGGCGC |
| seq17 | GCGCGATCCTACAGAATCGCGC | seq43 | GCGCCATGTGTTCTCGATGCGC |
| seq18 | GCGCAATCCGCCAAAGACGCGC | seq44 | GCGCCGATGACTAGGGTAGCGC |
| seq19 | GCGCAGACATTGCAGCAGGCGC | seq45 | GCGCGGTAGATACTCCCGGCGC |
| seq20 | GCGCGCAGGTCGAAATGGGCGC | seq46 | GCGCCCCGCTGTCAGACCGCGC |
| seq21 | GCGCATGGGACTCGGACAGCGC | seq47 | GCGCGACCCACCTCACGAGCGC |
| seq22 | GCGCGACAACATATTCCTGCGC | seq48 | GCGCACGATCTGCGGCAGGCGC |
| seq23 | GCGCTCCTCCGTCGGGTGGCGC | seq49 | GCGCGCAGTACATCACTTGCGC |
| seq24 | GCGCGGTGTGACCATAACGCGC | seq50 | GCGCACTTAACAATACGTGCGC |
| seq25 | GCGCTAACTTCGGCTCGTGCGC | seq51 | GCGCACGTGGACCGATAAGCGC |
| seq26 | GCGCTCGTTACGGGGGAAGCGC | seq52 | GCGCATAAAATATACGTAGCGC |

**Table S5.** Simulated oligomers comprising all 512 unique RNA pentamers.

| RNA | 5'-------------------3' | RNA | 5'-------------------3' |
|---|---|---|---|
| seq1 | GCGCAAAAUAGCCACUCUGCGC | seq27 | GCGCGGAAACGCUUUUUCGCGC |
| seq2 | GCGCCUCUCUUCAUAGUUGCGC | seq28 | GCGCUUUCUUGUGCAUACGCGC |
| seq3 | GCGCAGUUUUGCCAGAGGGCGC | seq29 | GCGCAUACCAACGGUGCUGCGC |
| seq4 | GCGCGAGGCACUGUUUACGCGC | seq30 | GCGCUGCUCUAUAUCCCUGCGC |
| seq5 | GCGCUUACUAAUCUCAUUGCGC | seq31 | GCGCCCCUGGGCUGGAAGGCGC |
| seq6 | GCGCCAUUAUACAAAUCAGCGC | seq32 | GCGCGAAGCUCCUGUGGUGCGC |
| seq7 | GCGCAUCAAUCGCAUCUUGCGC | seq33 | GCGCUGGUGAACGUCUAGGCGC |
| seq8 | GCGCUCUUAGAAGGGCAUGCGC | seq34 | GCGCCUAGCUGAAAGUCUGCGC |
| seq9 | GCGCGCAUUCAAGCCUGAGCGC | seq35 | GCGCGUCUCCAAGGACGCGCGC |
| seq10 | GCGCCUGAGCGCGGUUGAGCGC | seq36 | GCGCACGCCUAUCAUGCUGCGC |
| seq11 | GCGCUUGAGUUCGUCAAAGCGC | seq37 | GCGCUGCUUACCUAACCAGCGC |
| seq12 | GCGCCAAACCCCAAUUACGCGC | seq38 | GCGCACCAGUUGCGCCCGGCGC |
| seq13 | GCGCUUACACGCGAGGGGGCGC | seq39 | GCGCCCCGGAACCUUUAGGCGC |
| seq14 | GCGCGGGGCCGUGCGUAGGCGC | seq40 | GCGCUUAGCAAGUAGCGAGCGC |
| seq15 | GCGCGUAGUGUCGCCGGUGCGC | seq41 | GCGCGCGAAUUUAAUAAGGCGC |
| seq16 | GCGCCGGUACGACUGAUCGCGC | seq42 | GCGCUAAGGCCAUCCAUGGCGC |
| seq17 | GCGCGAUCCUACAGAAUCGCGC | seq43 | GCGCCAUGUGUUCUCGAUGCGC |
| seq18 | GCGCAAUCCGCCAAAGACGCGC | seq44 | GCGCCGAUGACUAGGGUAGCGC |
| seq19 | GCGCAGACAUUGCAGCAGGCGC | seq45 | GCGCGGUAGAUACUCCCGGCGC |
| seq20 | GCGCGCAGGUCGAAAUGGGCGC | seq46 | GCGCCCCGCUGUCAGACCGCGC |
| seq21 | GCGCAUGGGACUCGGACAGCGC | seq47 | GCGCGACCCACCUCACGAGCGC |
| seq22 | GCGCGACAACAUAUUCCUGCGC | seq48 | GCGCACGAUCUGCGGCAGGCGC |
| seq23 | GCGCUCCUCCGUCGGGUGGCGC | seq49 | GCGCGCAGUACAUCACUUGCGC |
| seq24 | GCGCGGUGUGACCAUAACGCGC | seq50 | GCGCACUUAACAAUACGUGCGC |
| seq25 | GCGCUAACUUCGGCUCGUGCGC | seq51 | GCGCACGUGGACCGAUAAGCGC |
| seq26 | GCGCUCGUUACGGGGGAAGCGC | seq52 | GCGCAUAAAAUAUACGUAGCGC |

**Table S6.** DNA and RNA sequences containing all tetramers (set14).

| DNA | 5'-------------------3' | RNA | 5'-------------------3' |
|---|---|---|---|
| seq1 | GCGCGGACGTTCAGCGACGCGC | seq1 | GCGCGGACGUUCAGCGACGCGC |
| seq2 | GCGCGCCCTGCATACAGTGCGC | seq2 | GCGCGCCCUGCAUACAGUGCGC |
| seq3 | GCGCTATATCTAGCTTGAGCGC | seq3 | GCGCUAUAUCUAGCUUGAGCGC |
| seq4 | GCGCGGAGTTGTCATGTTGCGC | seq4 | GCGCGGAGUUGUCAUGUUGCGC |
| seq5 | GCGCTCGAGCACTTAACCGCGC | seq5 | GCGCUCGAGCACUUAACCGCGC |
| seq6 | GCGCTGATCGGTGAGGATGCGC | seq6 | GCGCUGAUCGGUGAGGAUGCGC |
| seq7 | GCGCGACTACGAATGGTCGCGC | seq7 | GCGCGACUACGAAUGGUCGCGC |
| seq8 | GCGCGAGAAATTGCCTAAGCGC | seq8 | GCGCGAGAAAUUGCCUAAGCGC |
| seq9 | GCGCGCGCGGAAGGTAATGCGC | seq9 | GCGCGCGCGGAAGGUAAUGCGC |
| seq10 | GCGCGGCGTGGCCGGGAGGCGC | seq10 | GCGCGGCGUGGCCGGGAGGCGC |
| seq11 | GCGCGTACTGGGTCTTTAGCGC | seq11 | GCGCGUACUGGGUCUUUAGCGC |
| seq12 | GCGCTCCAAAACGGGGCTGCGC | seq12 | GCGCUCCAAAACGGGGCUGCGC |
| seq13 | GCGCAATAATCTGTGTCGGCGC | seq13 | GCGCAAUAAUCUGUGUCGGCGC |
| seq14 | GCGCATAGAGATGCGCGCGCGC | seq14 | GCGCAUAGAGAUGCGCGCGCGC |
